# Open database searching enables the identification and comparison of bacterial glycoproteomes without defining glycan compositions prior to searching

**DOI:** 10.1101/2020.04.21.052845

**Authors:** Ameera Raudah Ahmad Izaham, Nichollas E. Scott

**Affiliations:** Department of Microbiology and Immunology, University of Melbourne at the Peter Doherty Institute for Infection and Immunity, Melbourne 3000, Australia

## Abstract

Mass spectrometry has become an indispensable tool for the characterisation of glycosylation across biological systems. Our ability to generate rich fragmentation of glycopeptides has dramatically improved over the last decade yet our informatic approaches still lag behind. While glycoproteomic informatics approaches using glycan databases have attracted considerable attention, database independent approaches have not. This has significantly limited high throughput studies of unusual or atypical glycosylation events such as those observed in bacteria. As such, computational approaches to examine bacterial glycosylation and identify chemically diverse glycans are desperately needed. Here we describe the use of wide-tolerance (up to 2000 Da) open searching as a means to rapidly examine bacterial glycoproteomes. We benchmarked this approach using *N*-linked glycopeptides of *Campylobacter fetus subsp. fetus* as well as *O*-linked glycopeptides of *Acinetobacter baumannii* and *Burkholderia cenocepacia* revealing glycopeptides modified with a range of glycans can be readily identified without defining the glycan masses prior to database searching. Utilising this approach, we demonstrate how wide tolerance searching can be used to compare glycan utilisation across bacterial species by examining the glycoproteomes of eight Burkholderia species (*B. pseudomallei; B. multivorans; B. dolosa; B. humptydooensis; B. ubonensis, B. anthina; B. diffusa; B. pseudomultivorans*). Finally, we demonstrate how open searching enables the identification of low frequency glycoforms based on shared modified peptides sequences. Combined, these results show that open searching is a robust computational approach for the determination of glycan diversity within bacterial proteomes.

## INTRODUCTION

Protein glycosylation, the addition of carbohydrates to proteins, is a widespread and heterogeneous class of protein modifications [1-3]. Within Eukaryotes, multiple glycosylation systems have been identified [1–3] and up to 20% of the proteome is thought to be subjected to this class of modifications [4]. Within Eukaryotes, both *N*-linked and *O*-linked glycosylation systems are known to generate highly heterogeneous glycan structures [2, 3] with this glycan heterogeneity important for the function of glycoproteins [5, 6]. Although the glycan repertoire utilised in eukaryotic systems is thought to be large, the diversity within any given biological sample is constrained by the limited number of monosaccharides used in eukaryotic systems [7], as well as the expression of proteins required for the construction of glycans such as glycosyltransferases [8]. Experimentally, these constraints lead to only a limited number of glycans being produced across eukaryotic samples [9, 10] despite the large number of potential glycan structures [11, 12]. This limited diversity within both eukaryotic *N*-linked and *O*-linked glycans has enabled the development of glycan databases which have facilitated high throughput glycoproteomic studies [13] using tools such as Byonic [14] and pGlyco [15]. Unfortunately, these databases are not suitable for all glycosylation systems and fail to identify glycopeptides modified with novel or atypical glycans such as those found in bacterial glycosylation systems.

Within bacterial systems, glycosylation is increasingly recognised as a common modification [16–19]. While glycosylation in bacteria was first identified in the 1970s [20], it is only within the last two decades that it has become clear that this class of modifications is ubiquitous across bacterial genera [16, 18, 21]. Unlike eukaryotic systems, which utilise a relatively small set of monosaccharides, bacterial glycoproteins are decorated with a diverse range of monosaccharides [22] leading to a staggering array of glycan structures [23–32]. This glycan diversity represents a significant challenge to the field as it makes the identification of novel bacterial glycoproteins a non-trivial analytical undertaking. Yet, through advancements in mass spectrometry (MS) [28, 30, 33, 34], these once obscure modifications are increasingly recognisable and are now known to be essential for bacterial fitness [26, 35–38]. Despite our ability to generate rich bacterial glycopeptide data the field still largely uses manual interrogation to identify and characterise novel glycosylation systems [23–32]. This dependency on manual interrogation is not scalable, time-consuming and prone to human error, especially in the detection of glycoform heterogeneity. This is exemplified in our own experience characterising glycosylation in *Acinetobacter baumannii* where our initial analysis overlooked alternative methylated and deacetylated forms of glucuronic acid [26]. Thus, new approaches are needed to ensure bacterial glycosylation studies can be undertaken in a robust and high-throughput manner.

Wide precursor mass tolerance database searching, also known as ‘open’ or wildcard searching, is an increasingly popular approach for the detection of protein modifications within proteomic datasets [39–43]. The underlying premise of this approach is that by allowing a wide precursor mass tolerance, modified peptides can be detected by the difference in their observed mass from their expected mass. Importantly, this makes the identification of modifications independent of needing to define the modification in the initial search parameters. This approach has been utilised to examine chemical modifications such as formylation [44] and mis-alkylation [45] as well as large modifications such as DNA-peptide crosslinks [43] and eukaryotic glycosylation [46, 47]. Although this approach is effective, it is not without trade-offs being computationally more expensive than traditional searches leading to longer search times [48]. To date, these searches have typically been undertaken using **±** 500 Da tolerances [39–43] yet large delta mass windows of ±1000Da [43, 46] and even +3000Da [46, 47] have been reported. Despite the growing application of open database searching in eukaryotic proteomics, few bacterial studies have utilised this technique. This said, alternative strategies such as dependent peptide searching have been used in bacteria to track misincorporation of amino acids [49] and identify novel forms of glycosylation such as arginine-rhammnosylation [50].

In this study, we demonstrate that wide mass (up to 2000 Da) open database searching enables the rapid identification of bacterial glycopeptides without the need to assign glycan masses prior to database searching. Using Byonic™, which enables both glycopeptide and open database searching [14, 51], we benchmark wide mass open searching on three previously characterised bacterial glycosylation systems: the *N*-linked glycosylation system of *Campylobacter fetus subsp. fetus* NCTC10842 [25]; the *O*-linked glycosylation system of *Acinetobacter baumannii* ATCC17978 [26, 52, 53]; and the *O*-linked glycosylation system of *Burkholderia cenocepacia* J2315 [23, 37]. Each of these bacteria have increasingly complex proteomes (ranging from 1600 to nearly 7000 proteins) enabling us to assess the performance of open database searching across a range of proteome sizes. We find open database searching readily enabled previously characterised glycoforms and microheterogeneity to be identified across all samples. Applying this approach to representative species of the *Burkholderia genus* [23, 37], we provide the first snapshot of glycosylation across this genus. Consistent with the conservation of the biosynthetic pathway responsible for the Burkholderia *O*-linked glycans [23] all Burkholderia species examined predominately modify their glycoproteins with two glycan structures of similar composition. Excitingly, we demonstrate that open searching also enables low frequency glycoforms to be detected, highlighting that species-specific glycan structures do exist in Burkholderia. Thus, open database searching enables the identification of diverse bacterial glycan structures in a high-throughput manner.

## EXPERIMENTAL PROCEDURES

### Bacterial strains and growth conditions

*C. fetus subsp. fetus NCTC 10842* was grown on Brain-Heart Infusion medium (Hardy Diagnostics) with 5% defibrinated horse blood (Hemostat, Dixon, CA) under microaerobic conditions (10% CO_2_, 5% O_2_, 85% N_2_) at 37 °C as previously reported [25]. *Burkholderia pseudomallei* K96243 was grown as previously reported [54] in Luria Bertani (LB) broth. All other bacterial strains were grown overnight on LB agar at 37 °C as previously described [37]. Details on the strains, their origins, references and proteome databases used in this study are provided within Table 1.

**Table 1.** Strain list.

| Strains | Source (Description, Country, Year) | Reference | Proteome database |
| --- | --- | --- | --- |
| *C. fetus subsp. fetus* NCTC 10842 | Brain of sheep fetus, France, 1952 | [79] | Uniprot:<br>UP000001035 |
| *Acinetobacter baumannii* ATCC17978 | Fatal meningitis of a 4-month old infant, 1951 | [80] | GenBank assembly<br>accession:<br>GCA_001593425.2 |
| *Burkholderia pseudomallei* K96243 | Human clinical specimen, Thailand, 1996 | [81] | Uniprot:<br>UP000000605 |
| *Burkholderia cenocepacia* (LMG 16656 / J2315) | Human clinical specimen, United Kingdom, 1989 | [82] | Uniprot:<br>UP000001035 |
| *Burkholderia multivorans* MSMB2008 | Soil isolate, Australia, 2012 | [83] | *Burkholderia* Genome<br>Database [84],<br>Strain number: 3016 |
| *Burkholderia dolosa* AU0158 | Human clinical specimen, USA unknown | [85] | Uniprot database:<br>UP000032886 |
| *Burkholderia humptydooensis* MSMB43 | Water isolate, Australia, unknown | [83, 86] | *Burkholderia* Genome<br>Database [84],<br>Strain number: 4072 |
| *Burkholderia ubonensis*<br>MSMB22 | Soil isolate, Australia,<br>2001 | [85] | Burkholderia Genome<br>Database [84],<br>Strain number: 3404 |
| *Burkholderia anthina*<br>MSMB649 | Soil isolate, Australia,<br>2010 | [83] | Burkholderia Genome<br>Database [84],<br>Strain number: 2849 |
| *Burkholderia diffusa*<br>MSMB375 | Water isolate, Australia,<br>2008 | [83] | Burkholderia Genome<br>Database [84],<br>Strain number: 2966 |
| *Burkholderia pseudomultivorans*<br>MSMB2199 | Soil isolate, Australia,<br>2011 | [83] | Burkholderia Genome<br>Database [84],<br>Strain number: 3251 |

### Generation of bacterial lysates for glycoproteome analysis

Bacterial strains were grown to confluency on agar plates before being flooded with 5 mL of pre-chilled sterile phosphate-buffered saline (PBS) and bacterial cells collected by scraping. Cells were washed 3 times in PBS to remove media contaminates, then collected by centrifugation at 10,000 × *g* at 4°C for 10 min and then snap frozen. Snap frozen cell samples were resuspended in 4% SDS, 100mM Tris pH 8.0, 20mM Dithiothreitol (DTT) and boiled at 95°C with shaking at 2000rpm for 10 min. Samples were clarified by centrifugation at 17,000 × *g* for 10 min, the supernatants then collected, and protein concentration determined by a bicinchoninic acid assay (Thermo Scientific). 1mg of protein from each sample was acetone precipitated by mixing one volume of sample with 4 volumes of ice-cold acetone. Samples were precipitated overnight at −20°C and then spun down at 16,000 × g for 10 min at 0°C. The precipitated protein pellets were resuspended in 80% ice-cold acetone and precipitated for an additional 4 hours at −20°C. Samples were centrifuged at 17,000 × *g* for 10 min at 0°C, the supernatant discarded, and excess acetone driven off at 65°C for 5 min. Three biological replicates of each bacterial strain were prepared.

### Digestion of protein samples

Protein digestion was undertaken as previously described with minor alterations [28]. Briefly, dried protein pellets were resuspended in 6 M urea, 2 M thiourea in 40 mM NH_4_HCO_3_ then reduced for 1 hour with 20mM DTT followed by being alkylated with 40mM chloroacetamide for 1 hour in the dark. Samples were then digested with Lys-C (1/200 w/w) for 3 hours before being diluted with 5 volumes of 40 mM NH_4_HCO_3_ and digested overnight with trypsin (1/50 w/w). Digested samples were acidified to a final concentration of 0.5% formic acid and desalted with 50 mg tC18 Sep-Pak columns (Waters corporation, Milford, USA) according to the manufacturer’s instructions. tC18 Sep-Pak columns were conditioned with 10 bed volumes of Buffer B (0.1% formic acid, 80% acetonitrile), then equilibrated with 10 bed volumes of Buffer A* (0.1% TFA, 2% acetonitrile) before use. Samples were loaded on to equilibrated columns then columns washed with at least 10 bed volumes of Buffer A* before bound peptides were eluted with Buffer B. Eluted peptides were dried by vacuum centrifugation and stored at −20 °C.

### ZIC-HILIC enrichment of bacterial glycopeptides

ZIC-HILIC enrichment was performed as previously described with minor modifications [28]. ZIC-HILIC Stage-tips [55] were created by packing 0.5cm of 10 μm ZIC-HILIC resin (Millipore, Massachusetts, United States) into p200 tips containing a frit of C8 Empore™ (Sigma) material. Prior to use, the columns were washed with ultra-pure water, followed by 95% acetonitrile and then equilibrated with 80% acetonitrile and 5% formic acid. Digested proteome samples were resuspended in 80% acetonitrile and 5% formic acid. Samples were adjusted to a concentration of 3 μg/μL (a total of 300 μg of peptide used for each enrichment) then loaded onto equilibrated ZIC-HILIC columns. ZIC-HILIC columns were washed with 20 bed volumes of 80% acetonitrile, 5% formic acid to remove non-glycosylated peptides and bound peptides eluted with 10 bed volumes of ultra-pure water. Eluted peptides were dried by vacuum centrifugation and stored at −20 °C.

### Reverse phase LC-MS

ZIC-HILIC enriched samples were re-suspended in Buffer A* and separated using a two-column chromatography set up composed of a PepMap100 C18 20 mm × 75 μm trap and a PepMap C18 500 mm × 75 μm analytical column (Thermo Fisher Scientific). Samples were concentrated onto the trap column at 5 μL/min for 5 minutes with Buffer A (0.1% formic acid, 2% DMSO) and then infused into an Orbitrap Fusion™ Lumos™ Tribrid™ Mass Spectrometer (Thermo Fisher Scientific) at 300 nl/minute via the analytical column using a Dionex Ultimate 3000 UPLC (Thermo Fisher Scientific). 185-minute analytical runs were undertaken by altering the buffer composition from 2% Buffer B (0.1% formic acid, 77.9% acetonitrile, 2% DMSO) to 28% B over 150 minutes, then from 28% B to 40% B over 10 minutes, then from 40% B to 100% B over 2 minutes. The composition was held at 100% B for 3 minutes, and then dropped to 2% B over 5 minutes before being held at 2% B for another 15 minutes. The Lumos™ Mass Spectrometer was operated in a data-dependent mode automatically switching between the acquisition of a single Orbitrap MS scan (120,000 resolution) every 3 seconds and Orbitrap MS/MS HCD scans of precursors (NCE 30%, maximal injection time of 80 ms, AGC 1*10^5^ with a resolution of 15000). HexNAc oxonium ion (204.087 m/z) product-dependent MS/MS analysis [56] was used to trigger three additional scans of potential glycopeptides; a Orbitrap EThcD scan (NCE 15%, maximal injection time of 250 ms, AGC 2*10^5^ with a resolution of 30000); a ion trap CID scan (NCE 35%, maximal injection time of 40 ms, AGC 5*10^4^) and a stepped collision energy HCD scan (using NCE 32%, 40%, 48% for *N*-linked glycopeptide samples and NCE 28%, 38%, 48% for *O*-linked glycopeptide samples with a maximal injection time of 250 ms, AGC 2*10^5^ with a resolution of 30000). For *B. pseudomallei* K96243 glycopeptide enrichments, duplicate runs were undertaken as above with the Orbitrap EThcD scans modified to use the extended mass range setting (200 m/z to 3000 m/z) to improve the detection of high mass glycopeptide fragment ions [61].

### Data Analysis

Raw data files were batched processed using Byonic v3.5.3 (Protein Metrics Inc. [14]) with the proteome databases denoted within Table 1. Data was searched on a desktop with two 3.00GHz Intel Xeon Gold 6148 processors, a 2TB SDD and 128 GB of RAM using a maximum of 16 cores for a given search. For all searches, a semi-tryptic N-ragged specificity was set and a maximum of two missed cleavage events allowed. Carbamidomethyl was set as a fixed modification of cystine while oxidation of methionine was included as a variable modification. A maximum mass precursor tolerance of 5 ppm was allowed while a mass tolerance of up to 10 ppm was set for HCD fragments and 20 ppm for EThcD fragments. For open searches of *C. fetus fetus* samples (*N*-linked glycosylation), the wildcard parameter was enabled allowing a delta mass between 200 Da and 1600 Da on asparagine residues. For open searches of *O*-linked glycosylation samples, the wildcard parameter was enabled allowing a delta mass between 200 Da and 2000 Da on serine and threonine residues. For focused searches, all parameters listed above remained constant except wildcard searching which was disabled and specific glycoforms as identified from open searches included as variable modifications. To ensure high data quality, separate datasets from the same biological samples were combined using R and only glycopeptides with a Byonic score >300 were used for further analysis. This score cut-off is in line with previous reports highlighting that score thresholds greater than at least 150 are required for robust glycopeptide assignments with Byonic [44, 57]. It should be noted that a score threshold of above 300 resulted in false discovery rates of less than 1% for all combined datasets. Pearson correlation analysis of delta mass profiles was undertaken using Perseus [58]. Data visualization was undertaken using ggplot2 within R with all scripts included in the PRIDE uploaded datasets. All mass spectrometry proteomics data (Raw data files, Byonic search outputs, R Scripts and output tables) have been deposited into the PRIDE ProteomeXchange Consortium repository [59, 60] with the dataset identifier: PXD018587. Data can be accessed using the **username:**, **Password:** 7dcY3KOm

### Experimental Design and Statistical Rationale

For each bacterial strain examined three biological replicates were prepared and used for glycopeptide enrichments leading to three LC-MS runs per bacterial strain. Three separate enrichments were prepared and run to ensure an accurate representation of the observable glycoproteome. *B. pseudomallei* K96243 biological replicates were run twice with two different instrument methods to improve the characterisation of the 990 Da glycan. For *C. fetus fetus NCTC 10842* unenriched peptide samples were run with identical methods as those used for glycopeptide analysis to assess the presence of formylated glycans prior to enrichment.

## RESULTS

### Open database searching allows the identification of bacterial *N*-linked glycopeptides

Although open database searching enables the detection of a variety of modifications including eukaryotic glycosylation [46, 47], to our knowledge, it has not been applied to bacterial systems or the study of atypical forms of glycosylation. To enable the identification of glycopeptides with complex glycans, large delta mass windows are needed as even modest glycans (>three monosaccharides) would be larger than the 500 Da window typically used for open searching [39–43]. To enable wide mass searching we utilized Byonic which uses a peak look up approach for the identification of PSMs [14, 51]. This approach is both rapid and overcomes the combinatorial explosion which leads to long search times with database open searching [48, 51]. To assess the ability of Byonic based open searching to identify bacterial glycopeptides, we first examined glycopeptide enrichments of *C. fetus fetus NCTC 10842. C. fetus fetus* possesses a small proteome (~1600 proteins [61]) and is known to produce two *N*-linked glycans composed of β-GlcNAc-α1,3-[GlcNAc1,6-]GlcNAc-α1,4-GalNAc-α1,4-GalNAc-α1,3-diNAcBac (1243.507Da) and β-GlcNAc-α1,3-[Glc1,6-]GlcNAc-α1,4-GalNAc-α1,4-GalNAc-α1,4-diNAcBac (1202.481Da) where diNAcBac is the bacterial specific sugar 2,4-diacetamido-2,4,6 trideoxyglucopyranose [25].

We searched ZIC-HILIC glycopeptide enrichments of *C. fetus fetus* allowing a wildcard mass of 200Da to 1600 Da on asparagine enabling the processing of 3-hour LC-MS runs by open searching in less than 2 hours (Supplementary Figure 1A). Examination of the detected modifications by binning the observed delta masses in 0.001Da increments demonstrated a clear cluster of modifications with masses >1000Da (Figure 1A). Within these masses 1242.501Da and 1201.475Da were the most numerous delta masses observed (Figure 1A, Supplementary Table 1) yet these are one Dalton off the expected glycoforms of *C. fetus fetus* [25]. Close examination of the observed delta masses reveals evidence of mis-assignments of the mono-isotopic masses by the appearance of satellite peaks [42, 48] differing by exactly one Dalton (Supplementary Figure 2A). The mis-assignment of the mono-isotopic peak of large glycopeptides has been previously noted [62] and within Byonic is combated by allowing isotope re-assignment denoted as the “off-by-x” parameter. Examination of the “off-by-x” masses supports the inappropriate mass correction of the 1243/1202 Da glycans to the observed 1242/1201 delta masses (Supplementary Figure 2B). These results support that the 1243/1202 Da glycans are readily detected in *C. fetus fetus* samples using open database searching despite the splitting of the delta mass observations across multiple masses due to errors in mono-isotope assignments.

**Figure 1:**
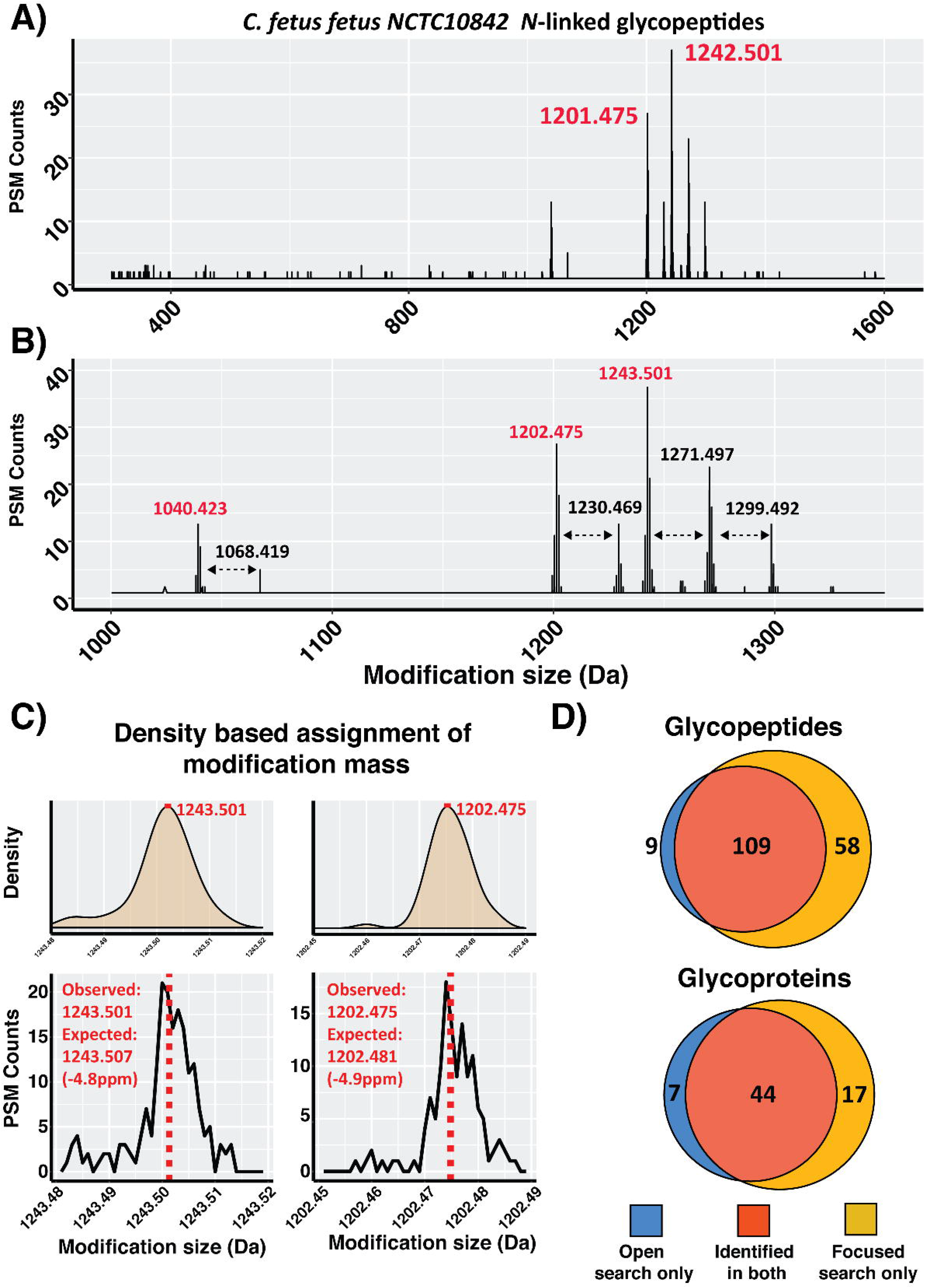
Open searching analysis of *C. fetus fetus* NCTC 10842 glycopeptides. **A)** *C. fetus fetus* glycopeptide delta mass plot of 0.001 Dalton increments showing the detection of PSMs modified with masses over 1000 Da. **B)** Zoomed view of the *C. fetus fetus* glycopeptide delta mass plot highlighting the most numerous observed delta masses; red masses correspond to non-formylated glycans while black masses correspond to formylated glycans. **C)** Density and zoomed delta mass plot of the *C. fetus fetus* glycan masses 1243.507Da and 1202.481Da. **D)** Comparison of the unique glycopeptide sequences and glycoproteins observed between open and focused searches across *C. fetus fetus* datasets.

Surprisingly, our open search also revealed additional glycoforms corresponding to formylated glycans (+27.99Da) as well as a modification corresponding to the loss of a HexNAc (−203.079Da) or Hex (−162.053Da) from the 1243Da or 1202Da glycans respectively (Figure 1B). MS/MS analysis supports these delta masses as unexpected but bona fide glycoforms (Supplementary figure 3A to J). Formylated glycans have been previously observed [25, 28] during ZIC-HILIC enrichment and are most likely artefacts due to the high concentrations of formic acid [44] used during enrichment. These formylated glycans represented a significant proportion of all potential glycopeptide PSMs (Figure 1B, Supplementary Table 1). Consistent with glycan formylation being artifactual, it is not observed on C. *fetus fetus* glycopeptides within unenriched samples (Supplementary figure 4). To assess the accuracy of the glycan masses obtained using open searching, we extracted the mean delta mass of the 1243 and 1202 Da glycans using a density based fitting approach [63] (Figure 1C and D). We find the open search defined mass of the 1243Da and 1202Da glycans are both within 5 ppm of the known masses [25] supporting that this approach allows high accuracy determination of large modifications. Finally, we assessed the proteome coverage of our open database approach to a traditional search using the seven identified glycoforms (1040.423Da, 1068.419Da, 1202.475Da, 1230.469Da, 1243.501Da, 1271.497Da, 1299.492Da, Figure 1B) as a focused search [64]. Consistent with previous studies focused searches outperformed open database searches [39, 46, 47] improving the identification of unique glycopeptides by 35% and glycoproteins by 28% (Figure 1D, Supplementary Table 2). This improvement was also associated with an increase in the average Byonic score from 456 to 491 for glycopeptide PSMs with identical MS/MS scans receiving an average 114 Byonic score increase between focused and open search assignments (Supplementary figure 5). These Focused searches also demonstrated that formylated glycans account for nearly a 1/3 of all glycopeptide assigned PSMs (>1246 formylated glycan PSMs out of the total 3824 glycopeptide PSMs, Supplementary Table 2). Combined, these results demonstrate open searching allows the detection of heterogeneous bacterial *N*-linked glycopeptides without the need to define glycans prior to searching.

### Open database searching allows the identification of bacterial *O*-linked glycopeptides

To assess open searching’s compatibility with bacterial O-linked glycopeptides, we examined glycopeptide enrichments of *A. baumannii ATCC17978*. The *A. baumannii* proteome is twice the size of *C. fetus fetus* (~3600 proteins [65]) with glycosylation of both serine and threonine residues reported to date [52]. Within this system, glycoproteins are modified predominantly with the glycan GlcNAc3NAcA4OAc-4-(β-GlcNAc-6-)-α-Gal-6-β-Glc-3-β-GalNAc where GlcNAc3NAcA4OAc corresponds to the bacterial sugar 2,3-diacetamido-2,3-dideoxy-α-D-glucuronic acid (glycan mass 1030.368 Da [52]). Importantly, this terminal glucuronic acid can also be found in methylated as well as un-acetylated states (corresponding to the glycan masses 1044.383 Da and 988.357 Da respectively [26, 52]). *A. baumannii* glycopeptide enrichments were searched allowing a wildcard mass of 200Da to 2000 Da on serine and threonine residues. The increased complexity of this search, both in terms of the number of amino acids potentially modified as well as the size of the proteome, resulted in a marked increase in the search times per data files to ~10 hours (Supplementary figure 1B). Within these samples, open searching readily enabled the identification of multiple delta masses of similar sizes to the expected glycans of *A. baumannii ATCC17978* as well as the unexpected glycoforms of 827.281 and 1058.358 Da (Figure 2A, Supplementary Table 3). These novel glycan masses are consistent with formylation (+27.99Da) as well as the loss of HexNAc (−203.079Da) from the 1030Da glycan with MS/MS analysis supporting these assignments (Supplementary Figure 6).

**Figure 2:**
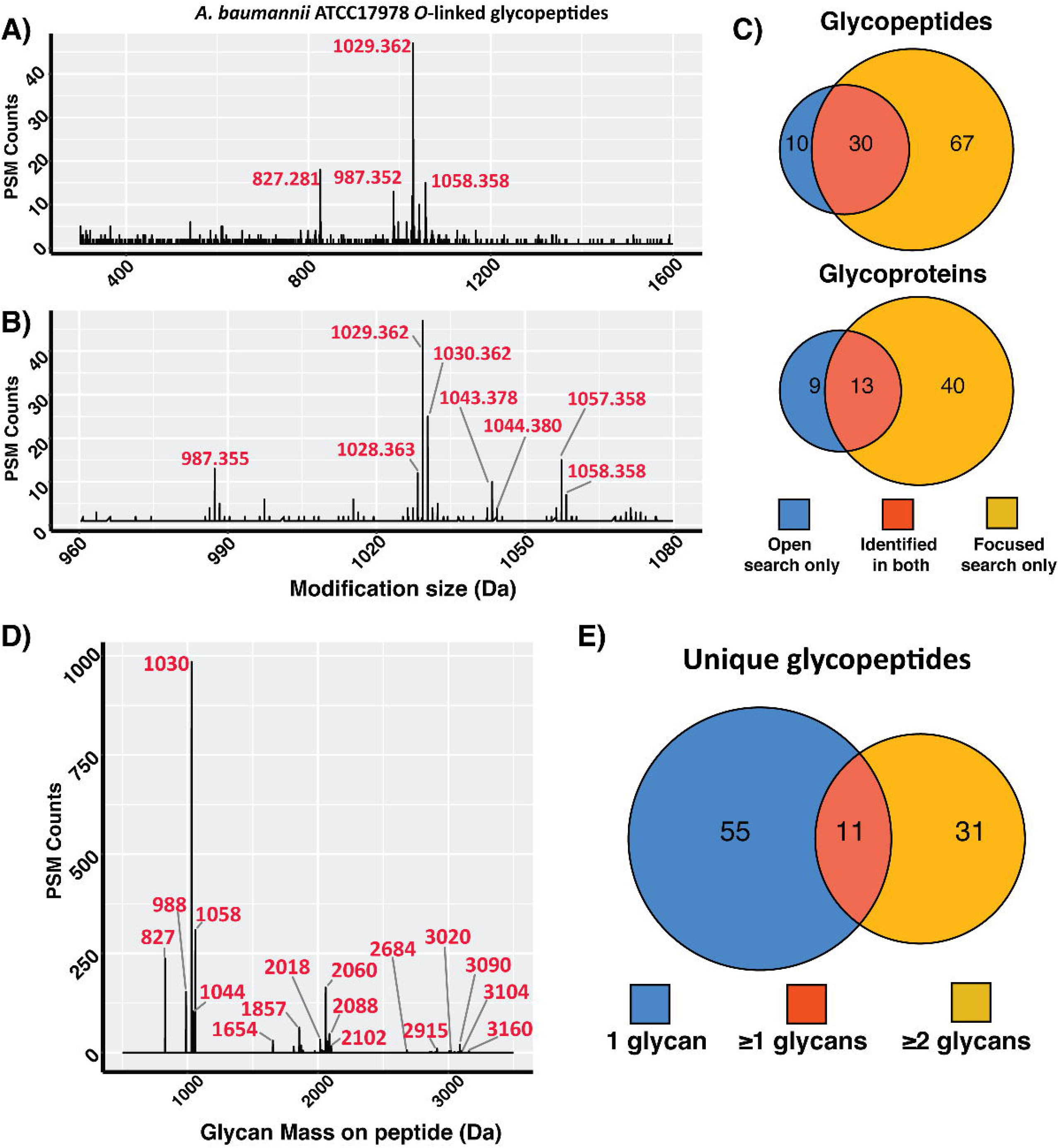
Open searching analysis of *A. baumannii ATCC17978* glycopeptides. **A)** *A. baumannii* glycopeptide delta mass plot of 0.001 Dalton increments showing the detection of PSMs modified with masses over 800 Da. **B)** Zoomed view of *A. baumannii* glycopeptide delta mass plot highlighting the most numerous observed modifications. **C)** Comparison of unique glycopeptide sequences and glycoproteins observed between open and focused searches across *A. baumannii* datasets **D)** Glycan mass plot showing the amount of glycan (in Da) observed on glycopeptides PSMs within the focused searches. **E)** Venn diagram showing the number of unique glycopeptide sequences grouped based on the number of glycans observed on these peptides.

Examination of these masses revealed the most numerous delta masses (1029.362 Da, 987.355 Da and 1043.378 Da) were one Dalton less than the expected *A. baumannii* glycan masses (Figure 2B [26, 52]). As with *C. fetus fetus,* inspection of these assignments reveals the incorrect application of the "off-by-x” parameter leading to the splitting of delta masses across multiple mass assignments separated by one Dalton (Supplementary figure 7). Using the masses 1030.368 Da, 988.357 Da, 1044.383 Da, 827.281 Da and 1058.358 Da, we researched these *A. baumannii* datasets to assess the performance of open to focused searching. In contrast to the ~35% increase in unique glycopeptides observed between open and focused searches in *C. fetus fetus* we noted a dramatic >240% improvement in unique glycopeptides identified within *A. baumannii* using focused searches (Figure 2C). To understand this dramatic improvement, we examined the 67 glycopeptides unique to the focused searches. Within these glycopeptides we noted a large proportion of PSMs corresponding to glycopeptides modified with multiple glycans (Figure 2D, Supplementary table 4). In fact, >20% (494 out of the total 2282 glycopeptide PSMs) corresponded to glycopeptides with greater than one glycan attached. Within these PSMs, 31 unique peptide sequences are only observed with >1 glycan attached (Figure 2E). The increased numbers of unique glycopeptides identified within focused searches were also associated with an increase in the mean Byonic scores as well as identical MS/MS scans receiving an average 149 Byonic score increase between focused and open search assignments (Supplementary Figure 8). Similar to *C. fetus fetus*, a large proportion of glycopeptide PSMs were identified with formylated glycans (>309 out of the total 2282 glycopeptide PSMs, Supplementary Table 4). It is important to note that the delta masses of multiply glycosylated peptides fall outside the 2000 Da window used for open searching making the inability to detect these glycopeptides an expected limitation of the search parameters. Thus, although open searching enables the rapid identification of glycopeptides, large glycans / multiply glycosylated peptides can be overlooked supporting the value of a two-step (open followed by focused) searching approach.

### Open database searching enables the identification of glycosylation within large proteomes

As open searching enabled the identification of both N and O-linked glycopeptides, we sought to explore the compatibility of this approach with larger proteomes using glycopeptide enriched samples from the bacteria *B. Cenocepacia* J2315 as a model. The *B. Cenocepacia* proteome encodes ~7000 proteins [66] and possesses an *O*-linked glycosylation system responsible for modifying at least 23 proteins [37]. Previously, we showed that this glycosylation system transfers two glycans composed of β-Gal-(1,3)-α-GalNAc-(1,3)-β-GalNAc and Suc-β-Gal-(1,3)-α-GalNAc-(1,3)-β-GalNAc where Suc is Succinyl with these glycans corresponding to the masses 568.211Da and 668.228Da respectively [23, 37]. As with *A. baumannii,* the increased complexity of this proteome led to an increase in the search time with individual data files taking ~20hours to process (Supplementary figure 1C). These open searches revealed the presence of the expected glycoforms of *B. cenocepacia* (568.207Da and 668.223Da) as well as additional formylated variants (596.202Da, 624.197, and 696.218Da) leading to the identification of five unique glycoforms (Figure 3A, Supplementary Table 5). Unlike the large glycans of *C. fetus fetus and A. baumannii,* it is notable that the mono-isotopic mass of the known *B. Cenocepacia* glycans [37] were correctly assigned (Figure 3A). Thus, this supports that for smaller glycans mis-assignment of the mono-isotopic masses during open searches does not appear as problematic.

**Figure 3:**
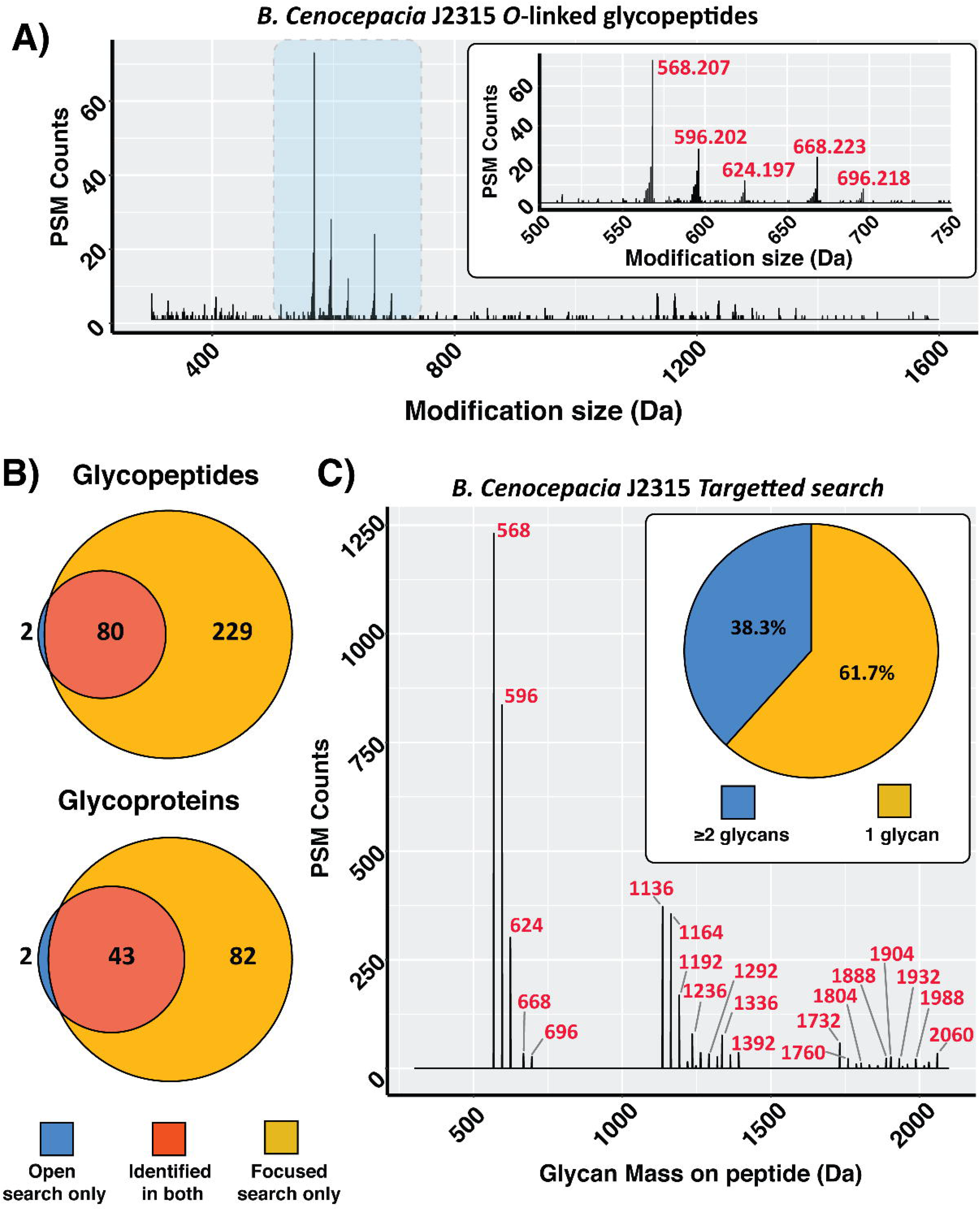
Open searching analysis of *B. Cenocepacia* J2315 glycopeptides. **A)** *B. Cenocepacia* glycopeptide delta mass plot of 0.001 Dalton increments showing the detection of PSMs modified with masses over 500 Da. Highlighted area shown in zoomed panel. **B)** Comparison of the unique glycopeptide sequences and glycoproteins observed between open and focused searches across *B. Cenocepacia* datasets **C)** Glycan mass plot showing the amount of glycan (in Da) observed on glycopeptides PSMs within the focused searches. Nearly 40% of all glycopeptide PSMs are decorated with two or more glycans.

Incorporating these glycoforms into focused searches again led to a dramatic ~4-fold increase in the number of glycopeptides and a ~2-fold increase in the total number glycoproteins identified compared to open searches (Figure 3B, Supplementary Table 6). Although the improvement in the total number of identifications was associated with a decrease in the mean Byonic score (from 728 to 700) the assigned scores of identical MS/MS scans reveals focused searches lead to an increase in the average Byonic score of 175 compared to assignments from open searches (Supplementary Figure 9). As the dramatic improvement in the glycoproteome coverage of *A. baumannii* was partially driven by the detection of multiply glycosylated peptides we examined the amount of glycosylation within glycopeptide PSMs in *B. Cenocepacia*. As *B. Cenocepacia* glycopeptides modified with multiple glycans would be less than 2000 Da, we were surprised by the limited number of multiply glycosylated peptides identified within our open searches (<10% of all identified glycopeptides, Supplementary figure 10). In contrast, focused searches identified ~40% of all PSMs (Figure 3C, 1508 out of 3937 identified glycopeptide PSMs) corresponded to multiply modified peptides. As with *C. fetus fetus* and *A. baumannii* formylated glycans make up nearly 50% of all glycopeptide PSMs (Figure 3C, Supplementary table 6). This data supports that, although open searching performs well for singly modified peptides, this approach appears to underrepresent multiply glycosylated peptides even if the combined mass of the glycan is within the range of the open search.

### Open database searching allows the screening of glycan utilisation across biological samples

Having established that open searching enables the identification of a range of glycans, we sought to explore if this could also facilitate the comparison of glycan diversity across bacterial samples. Recently, we reported that a single-loci was responsible for the generation of the O-linked glycans in *B. Cenocepacia* and that this loci is conserved across the *Burkholderia* [23]. Although these results support that Burkholderia species utilise similar glycans, it has been noted that within other bacterial genera extensive glycan heterogeneity exists [25, 26, 29, 32]. As glycan heterogeneity can be challenging to predict, we reasoned that open searching would provide a means to assess the similarities in glycans used across Burkholderia species. We examined glycopeptide enrichments from eight species of Burkholderia (*B. pseudomallei K96243; B. multivorans MSMB2008; B. dolosa AU0158; B. humptydooensis MSMB43; B. ubonensis MSMB22, B. anthina MSMB649; B. diffusa MSMB375; and B. pseudomultivorans MSMB2199*). Examination of the delta masses observed across these eight species demonstrates that the 568Da and 668Da glycoforms as well as their formylated variants are present in all strains (Figure 4A and Supplementary Figure 11, Supplementary Tables 7 to 14). Having generated “delta mass fingerprints” for each species, we assessed if these profiles could enable the comparison of glycan utilisation using Pearson correlation and hierarchical clustering (Figure 4B and Supplementary Figure 12). Consistent with the similarities in the delta mass fingerprints Pearson correlation and hierarchical clustering resulted in the grouping of all Burkholderia species compared to the delta mass fingerprints of *C. fetus fetus* and *A. baumannii* (Figure 4B). These results support that consistent with the conservation of the glycosylation loci within Burkholderia, the major glycoforms observed within Burkholderia species, based on mass at least, are identical. It should be noted that as with the above glycopeptide datasets, focused searches significantly improved the identification of glycopeptides and glycoproteins in all Burkholderia species (Supplementary figure 13, Supplementary Table 15 to 22).

**Figure 4:**
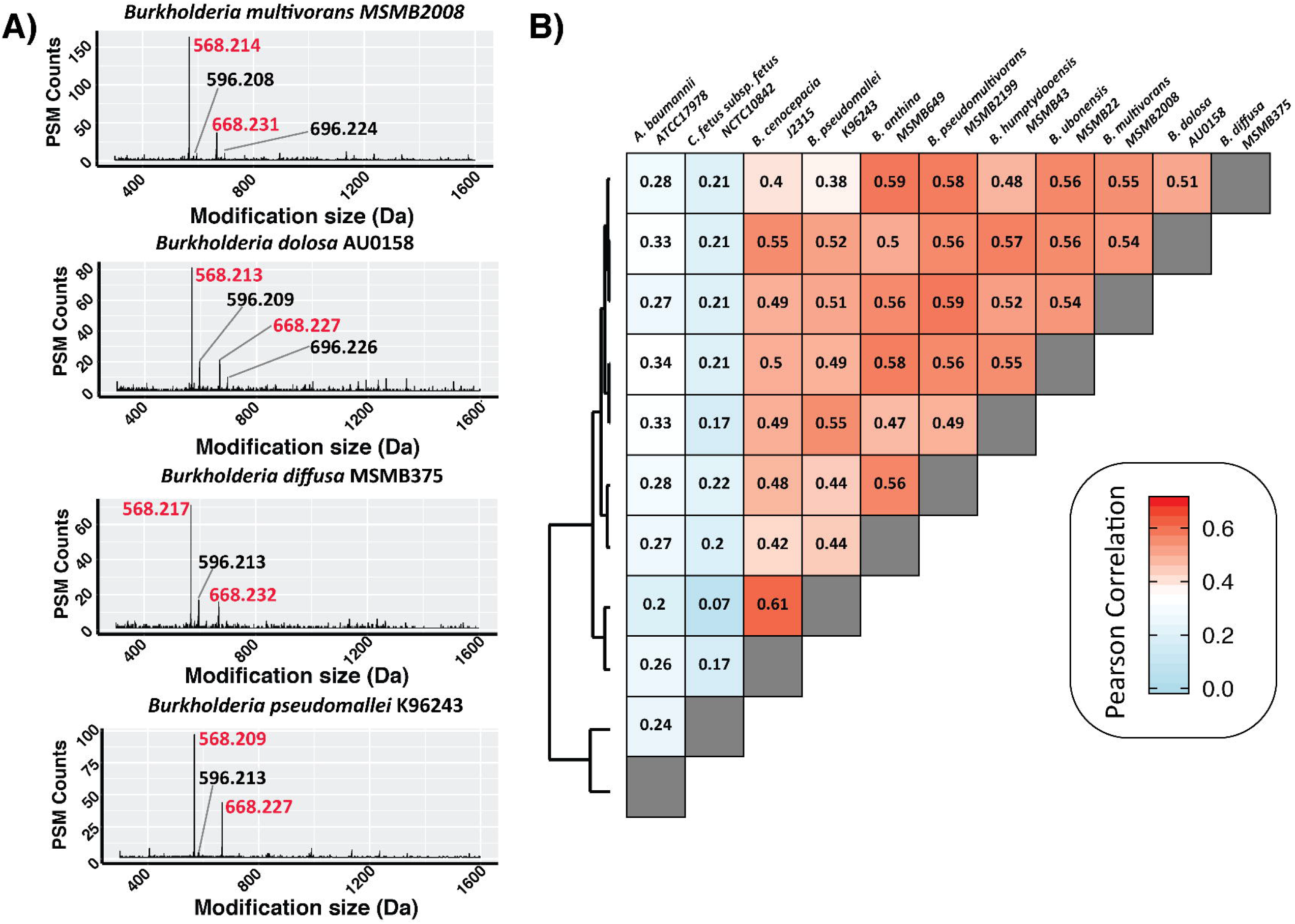
Comparison of Burkholderia glycoproteomes using open searching. **A)** Representative delta mass plots of four out of the eight Burkholderia strains examined demonstrating the 568Da and 668Da glycans are frequently identified delta masses in Burkholderia glycopeptide enrichments. Formylated glycans are denoted in black while Burkholderia O-linked glycans are in red. **B)** Pearson correlation and clustering analysis of delta mass plots enable the comparison and grouping of samples.

### Open database searching allows the detection of glycoforms identified at a low frequency based on known glycosylatable peptides

In addition to allowing the comparison of glycan diversity across species, we reasoned that open searching would also allow the identification of novel glycans based on the shared utilization of glycosylatable peptide sequences. Within bacterial glycosylation studies, proteins compatible with different glycosylation machinery are routinely used to “fish” out glycans used for protein glycosylation in different bacterial species [26, 32]. Similarly, by focusing on peptides modified with the 568/668Da glycans we hypothesized this would provide the means to detect alternative glycans used for glycosylation within Burkholderia species. To assess this, we examined the glycopeptide enrichments of *B. pseudomallei* K96243 filtering for delta masses only observed on peptide sequences also modified with either the 568/668Da glycans (Figure 5A). Examination of these delta masses readily revealed the presence of PSMs matching the modification of peptides with single (203.077Da) or double (406.158Da) HexNAc residues, two 568Da glycans (1136.422Da) and an unexpected mass at 990.390Da (Figure 5A). Examination of PSMs assigned to this 990Da delta mass revealed a linear glycan composed of HexNAc-Heptose-Heptose-188-215 where the 188 Da and 215 Da are moieties of unknown composition (Figure 5B). Incorporation of this unexpected glycan mass into a focused search with the known Burkholderia glycans demonstrated that the 990.390Da glycan is observed on multiple peptide substrates yet less than 6% of all glycopeptide PSMs correspond to this novel glycan (Figure 5C). Thus, this demonstrates that open searching provides an effective means to detect unexpected glycoforms which could be overlooked due to the low frequency of their occurrence in glycoproteomic datasets.

**Figure 5:**
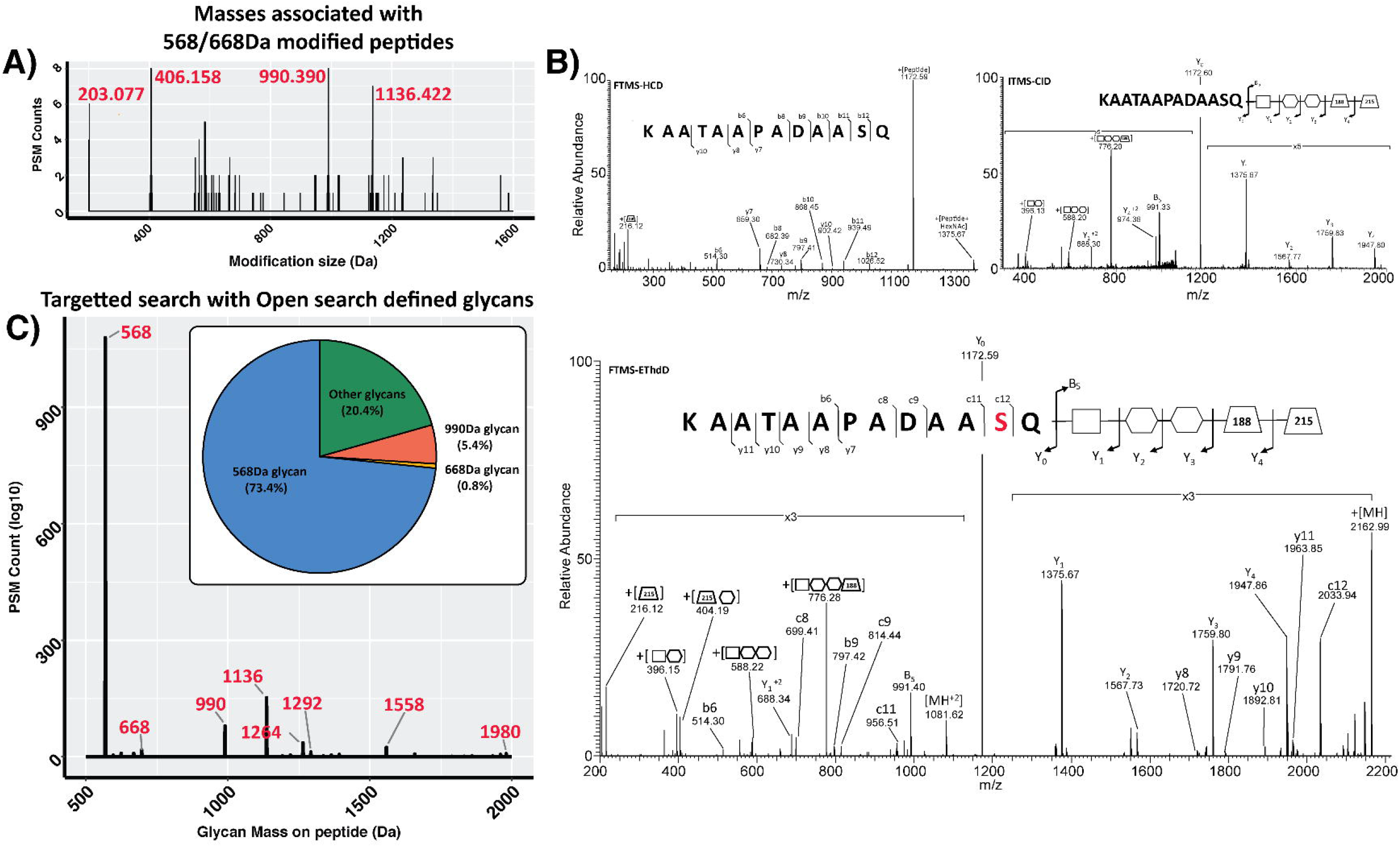
Identification of minor glycoforms within *B. pseudomallei* K96243. **A)** Delta mass plot, binned by 0.001 Dalton increments, showing delta masses observed for peptide sequences also modified with the 568 or 668D glycans. **B)** MS/MS analysis (FTMS-HCD, ITMS-CID and FTMS-EThcD) supporting the assignment of a linear glycan of HexNAc-Heptose-Heptose-188-215 attached to the peptide KAATAAPADAASQ. **C)** Glycan mass plot showing the amount of glycan (in Da) observed on glycopeptides PSMs within focused searches. Only ~6% of all PSMs observed are modified with the 990 Da glycan.

## DISCUSSION

MS analysis of glycoproteomic samples typically requires knowledge of both the proteome and possible glycan compositions to facilitate software-based identification [13]. As bacterial glycosylation systems do not utilize glycans found in eukaryotic glycan databases [23–32], we sought to establish an alternative approach for the high-throughput analysis of bacterial glycoproteomes. Within this work we demonstrate that wide mass open database searching enables the identification of bacterial glycosylation without the need to define glycan masses prior to searching. This approach overcomes a significant bottleneck in the identification and characterization of novel bacterial glycosylation systems. We demonstrate that a range of diverse glycan structures, both reported [25, 26, 37, 52] and unreported, such as the 990 Da glycan observed in *B. pseudomallei* K96243, as well as glycan artefacts such as formylated glycans can be identified using this approach. In addition, we also demonstrate that open database searches can be used to provide a simple means to compare delta mass profiles across biological samples. This enables a straightforward method to compare and contrast bacterial glycoproteomes, enabling the grouping of Burkholderia profiles from non-similar glycan profiles such as those seen in *C. fetus fetus* or *A. baumannii*.

Within this work we utilized open searching within Byonic, a widely used tool in the glycoproteomic community for the analysis of glycosylation [57, 67, 68]. This enabled us to directly compare the performance of open searches to focused glycopeptide searches within the same platform. We observed a marked improvement in glycopeptide and glycoprotein identifications within focused searches, especially for glycopeptides modified with multiple glycans. As a number of unique considerations are needed for optimal glycopeptide identification, such as accounting for glycan fragments [13, 69], this improvement in performance is unsurprising. Consistent with this we observed an increase in the Byonic scores within focused searches compared to open searches for most datasets (Supplementary figure 5, 8 and 9). This improvement translates to an increase in the numbers of unique glycopeptides and glycoproteins identified by ~35% to 240%. This is in line with previous studies [46, 47] and the observation that non-optimized glycopeptide analysis can lead to a reduction in glycoproteome coverage [64]. Although Byonic was used within this study it should be noted alternative non-commercial platforms such as MSfragger [43] also allow open searching. In our hands MSfragger performed comparably to Byonic for the identification of glycoforms using open searches (Supplementary Figure 14A to F). Yet, as with our open Byonic searches MSfragger did not identify as many unique glycopeptides / glycoproteins as focused Byonic searches (Supplementary Figure 15A to F). These results demonstrate that open searching can be used to identify glycopeptides, yet due to the unique challenges associated with glycopeptide identification open searches can be less sensitive than focused searches.

Although we have utilised open searching with masses up to 2000 Da wider mass ranges can and have been used to identify glycopeptides. In fact, for eukaryotic glycosylation analysis open searches with +3000 Da have been demonstrated enabling the identification of glycans of up to ~2000 Da in size [46, 47]. Although multiple open searching tools allow the maximum delta mass range to be widened this can be at the expense of search times [48]. It should be noted for open searching there is likely an upper limit for the maximum delta mass which can be used before no additional high confidence assignments are gained. This maximum limit is not only determined by the analytes being examined but also the MS/MS acquisition parameters. For example, for glycopeptides as the amount of glycan decorating peptides increases the optimal amounts of collision energy and ETD reaction times begin to diverge from standard parameters [70, 71]. Thus, even using open searching some specific subsets of peptides may still be unidentifiable without careful tailoring of the data acquisition methods.

At its core, this analytical approach utilizes a “strength in numbers” based strategy for the detection of glycans. A key strength of this approach is that it does not require the identification of unmodified versions of a peptide for the modified forms to be identified as required in dependent peptide based approaches [49, 50]. This independence of the need for unmodified peptides makes this approach compatible with enrichment strategies such as ZIC-HILIC glycopeptide enrichment. This is important as for optimal performance this approach requires large numbers of PSMs with identical delta masses. It should be noted that within ZIC-HILIC enrichments bacterial glycopeptides are still only a minor proportion of observed peptides being <10% of the identified PSMs (Supplementary Figure 16). Within this work we focused on bacterial glycosylation systems known to target multiple protein substrates [16, 18], ensuring large numbers of unique PSMs / peptide sequences would be identified. We found that, within glycopeptide enrichments, the known glycoforms of *C. fetus fetus*, *A. baumannii* and *B. cenocepacia* were easily detected while infrequently observed glycans, such as the 990Da glycan observed within *B. pseudomallei*, required additional filtering to distinguish this from background signals. This supports that although open searching enables the detection of glycoforms, it is sensitive to the frequency at which modification events are observed within datasets. Although we utilized filtering based on glycosylatable peptide sequences to identify low frequency events, recently Kernel density estimation based fitting approaches were shown to effectively address this issue in a more general manner [63]. Thus, open database searching provides multiple approaches to identify modifications even those which are poorly resolved from background.

A surprising observation within our open searches was the commonality of glycan formylation events within enriched glycopeptide samples. Recently it was shown that formic acid concentrations as low as 0.1% could lead to peptide formylation [44]. For the enrichment of bacterial glycopeptides ZIC-HILIC enrichment with 5% formic acid / 80% acetonitrile has been extensively used [24, 26-28, 36, 37, 52, 72] yet the observation of formylation artefacts on a substantial number of glycopeptide PSMs (Supplementary Table 2, 4, 6, 15 to 22) raises concerns about this protocol. Multiple reagents including Tris [73] and alkaline solutions such as ammonium hydroxide / sodium hydroxide [74] can lead to alterations within glycan structures necessitating the judicious use of these chemicals within glycan/glycopeptide sample preparation protocols. While 5% formic acid improves the selectivity for glycopeptides within ZIC-HILIC enrichments [75] alternative acids can also be used. Previously Mysling *et al* demonstrated that formic acid could be substituted for TFA to improve the enrichment of glycopeptides [76], while Ding *et al* noted that hydrochloric acid was an effective ion-pairing agent for normal phase enrichment of bacterial glycopeptides [77]. Thus, the observation of formylation highlights that alternative buffers should be considered for future glycopeptide studies and that formylated glycans need to be considered when analyzing glycopeptide datasets where formic acid has been used during glycopeptide enrichments.

Although open searching enabled the identification of glycosylation within all bacterial samples, the analysis of *C. fetus fetus* and *A. baumannii* datasets highlighted the commonality at which mono-isotopic masses of glycopeptides with large glycans (>1000 Da) are mis-assigned. This problem has been highlighted previously [62] and is not unique to glycopeptides with the mono-isotopic mass of other large biomolecules such as cross-linked peptide shown to be mis-assigned in 50 to 75% of PSMs [78]. This mis-assignment of mono-isotopic masses leads to the splitting of the number of observed PSMs with a specific glycan mass across multiple mass channels. Although we demonstrate that these mis-assigned glycopeptides can be readily identified by examining the “off-by-x” parameter, it should be noted that this splitting dilutes the observable glycopeptides at a specific mass, complicating the analysis of glycoproteomes from open searches. This complication, coupled with the lower sensitivity of glycopeptide identification with open searching compared to focused searches discussed above, supports that open searching is a useful discovery tool yet typically under reports unique glycopeptides and glycoproteins within datasets. The simplest solution to this issue is to use open searching as a means to identify glycans which can then be included as variables modification within a focused search. As highlighted above, this significantly improves glycoproteome coverage and in our hands provided the flexibility of being able to detect novel glycans yet also ensured optimal identification of glycopeptides. Automated pipelines using multi-step searching have already been demonstrated [42, 48] yet to our knowledge these have not been optimized or implemented for glycopeptide identification. Thus, we recommend a multi-step analysis to enable the identification of atypical glycosylation, using open searching to define glycans which are then incorporated into focused searches.

Finally, it should be noted that although not the subject of this manuscript, the glycoproteins/glycopeptides identified in this work are themselves a useful resource for the bacterial glycosylation community. Previous studies on *C. fetus fetus*, *A. baumannii* and *B. cenocepacia* identified a total of 26, 26 and 23 unique glycoproteins respectively [25, 26, 37] yet the majority of these studies were undertaken on previous generations of MS instrumentation. Within this work, undertaken on a current generation instrument, we observed a marked improvement in the number of glycoproteins identified with 61 (2.3-fold), 53 (2.0-fold) and 125 (5.4-fold) glycoproteins identified in *C. fetus fetus*, *A. baumannii* and *B. cenocepacia* respectively. Similarly, our glycoproteomic analysis of the 8 Burkholderia species highlights that at least 70 proteins are glycosylated within each Burkholderia species. Taken together, this work highlights that the glycoproteomes of most bacterial species are likely far larger than earlier studies suggested with open searching providing an accessible starting point to probe these systems.

## ABBREVIATIONS

MS: mass spectrometry
LB: Luria Bertani
PBS: phosphate-buffered saline
SDS: sodium dodecyl sulfate
NCE: normalized collisional energy
ZIC-HILIC: Zwitterionic Hydrophilic Interaction Liquid Chromatography
Tris: Tris(hydroxymethyl)aminomethane
TFA: Trifluoroacetic acid
DTT: Dithiothreitol
EThcD: Electron-transfer/higher-energy collision dissociation
HCD: higher-energy collision dissociation
CID: Collision-induced dissociation
AGC: Automatic Gain Control
DMSO: Dimethyl sulfoxide
diNAcBac: 2,4-diacetamido-2,4,6 trideoxyglucopyranose
PSMs: peptide spectrum matches
Glc: Glucose
Gal: galactose
GalNAc: N-Acetylgalactosamine
GlcNAc: N-Acetylglucosamine
HexNAc: N-acetylhexoseamine
Hex: Hexose
2,3-diacetamido-2,3-dideoxy-α-D-glucuronic acid: GlcNAc3NAcA4OAc

## ACKNOWLEDGEMENTS

This work was supported by a National Health and Medical Research Council of Australia (NHMRC) project grant awarded to NES (APP1100164). We thank the Melbourne Mass Spectrometry and Proteomics Facility of The Bio21 Molecular Science and Biotechnology Institute for access to MS instrumentation and Byonic. We would like to thank Christine Szymanski and Justin Duma for the kind gift of *C. fetus fetus* NCTC 10842 lysates; Mitali Sarkar-Tyson and Nicole Bzdyl for providing the *B. pseudomallei* K96243 lysates; Deborah Yoder-Himes, Mark Mayo, Bart Currie and Amy Cain for kindly providing Burkholderia strains for this analysis as well as Chris McDevitt and Saleh Alquethamy for providing A. baumannii ATCC17978. We thank Ben Parker and Nick Williamson for their critical feedback on the manuscript.

## DATA AVAILABILITY

All raw data is available through the PRIDE repository, PRIDE accession: PXD018587.

**username:**, **Password:** 7dcY3KOm

## Notes

### Competing Interest Statement

The authors have declared no competing interest.

### Summary of Updates

This manuscript has been updated to address reviewers comments. These have greatly improved the readability as well as highlighted important missed citations which have now been included in this version of the manuscript.

