## Supplementary figures and supplementary table legends for "Open database searching enables the identification and comparison of bacterial glycoproteomes without defining glycan compositions prior to searching"

**Key words:** Glycosylation, Post-translational modifications, Proteomics, wide tolerant database searching, Open searching, Glycopeptides, bacterial glycosylation, atypical glycosylation

**SUPPLEMENTARY TABLES**

**Supplementary table 1: Open searching results for *C. fetus fetus*.** The combined Byonic searches results of three biological replicates of *C. fetus fetus* showing all modified peptides with a Score over 300.

**Supplementary table 2: Focused database results for *C. fetus fetus***. The combined Byonic searches results of three biological replicates of *C. fetus fetus* showing all modified peptides with a Score over 300.

**Supplementary table 3: Open searching results for *A. baumannii*.** The combined Byonic searches results of three biological replicates of *A. baumannii* showing all modified peptides with a Score over 300.

**Supplementary table 4: Focused database results for *A. baumannii***. The combined Byonic searches results of three biological replicates of *A. baumannii* showing all modified peptides with a Score over 300.

**Supplementary table 5: Open searching results for *B. Cenocepacia*.** The combined Byonic searches results of three biological replicates of *B. Cenocepacia* showing all modified peptides with a Score over 300.

**Supplementary table 6: Focused database results for *B. Cenocepacia***. The combined Byonic searches results of three biological replicates of *B. Cenocepacia* showing all modified peptides with a Score over 300.

**Supplementary table 7: Open searching results for *B. pseudomallei*.** The combined Byonic searches results of three biological replicates of *B. pseudomallei* showing all modified peptides with a Score over 300.

**Supplementary table 8: Open searching results for *B. diffusa*.** The combined Byonic searches results of three biological replicates of *B. diffusa* showing all modified peptides with a Score over 300.

**Supplementary table 9: Open searching results for *B. dolosa*.** The combined Byonic searches results of three biological replicates of *B. dolosa* showing all modified peptides with a Score over 300.

**Supplementary table 10: Open searching results for *B. multivorans*.** The combined Byonic searches results of three biological replicates of *B. multivorans* showing all modified peptides with a Score over 300.

**Supplementary table 11: Open searching results for *B. ubonensis*.** The combined Byonic searches results of three biological replicates of *B. ubonensis* showing all modified peptides with a Score over 300.

**Supplementary table 12: Open searching results for *B. pseudomultivorans*.** The combined Byonic searches results of three biological replicates of *B. pseudomultivorans* showing all modified peptides with a Score over 300.

**Supplementary table 13: Open searching results for *B. anthina*.** The combined Byonic searches results of three biological replicates of *B. anthina* showing all modified peptides with a Score over 300.

**Supplementary table 14: Open searching results for *B. humptydooensis*.** The combined Byonic searches results of three biological replicates of *B. humptydooensis* showing all modified peptides with a Score over 300.

**Supplementary table 15: Focused database results for *B. pseudomallei*.** The combined Byonic searches results of three biological replicates of *B. pseudomallei* showing all modified peptides with a Score over 300.

**Supplementary table 16: Focused database results for *B. diffusa*.** The combined Byonic searches results of three biological replicates of *B. diffusa* showing all modified peptides with a Score over 300.

**Supplementary table 17: Focused database results for *B. dolosa*.** The combined Byonic searches results of three biological replicates of *B. dolosa* showing all modified peptides with a Score over 300.

**Supplementary table 18: Focused database results for *B. multivorans*.** The combined Byonic searches results of three biological replicates of *B. multivorans* showing all modified peptides with a Score over 300.

**Supplementary table 19: Focused database results for *B. ubonensis*.** The combined Byonic searches results of three biological replicates of *B. ubonensis* showing all modified peptides with a Score over 300.

**Supplementary table 20: Focused database results for *B. pseudomultivorans*.** The combined Byonic searches results of three biological replicates of *B. pseudomultivorans* showing all modified peptides with a Score over 300.

**Supplementary table 21: Focused database results for *B. anthina*.** The combined Byonic searches results of three biological replicates of *B. anthina* showing all modified peptides with a Score over 300.

**Supplementary table 22: Focused database results for *B. humptydooensis*.** The combined Byonic searches results of three biological replicates of *B. humptydooensis* showing all modified peptides with a Score over 300.

**SUPPLEMENTARY FIGURES**

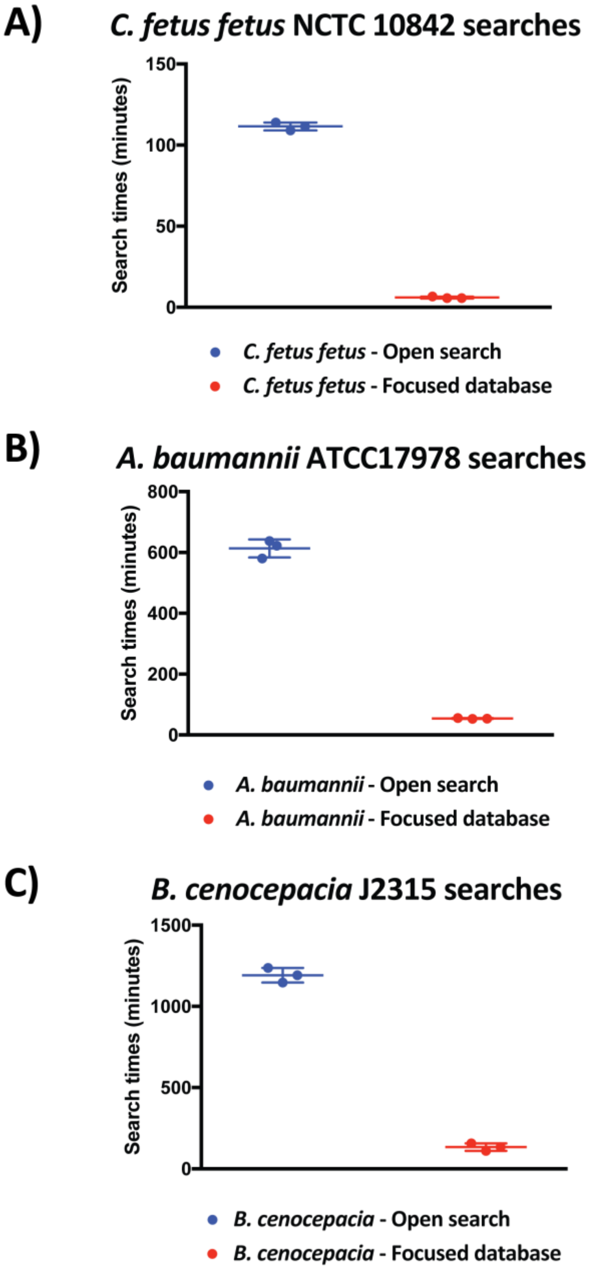


**Supplementary Figure 1: Comparison of search times with open and focused searches.** Search times for the open and focused searches are provided for *C. fetus fetus*, *A. baumannii* and *B. cenocepacia*. With increasing complexity of the proteome and search parameters we noted an increase in search times. The type of glycosylation and proteome size for these samples are: **A)** *C. fetus fetus*, *N*-linked glycosylation searched against a ~1600 protein proteome; **B)** *A. baumannii* *O*-linked glycosylation searched against a ~3600 protein proteome; and **C)** *B. cenocepacia* *O*-linked glycosylation searched against a ~7000 protein proteome.

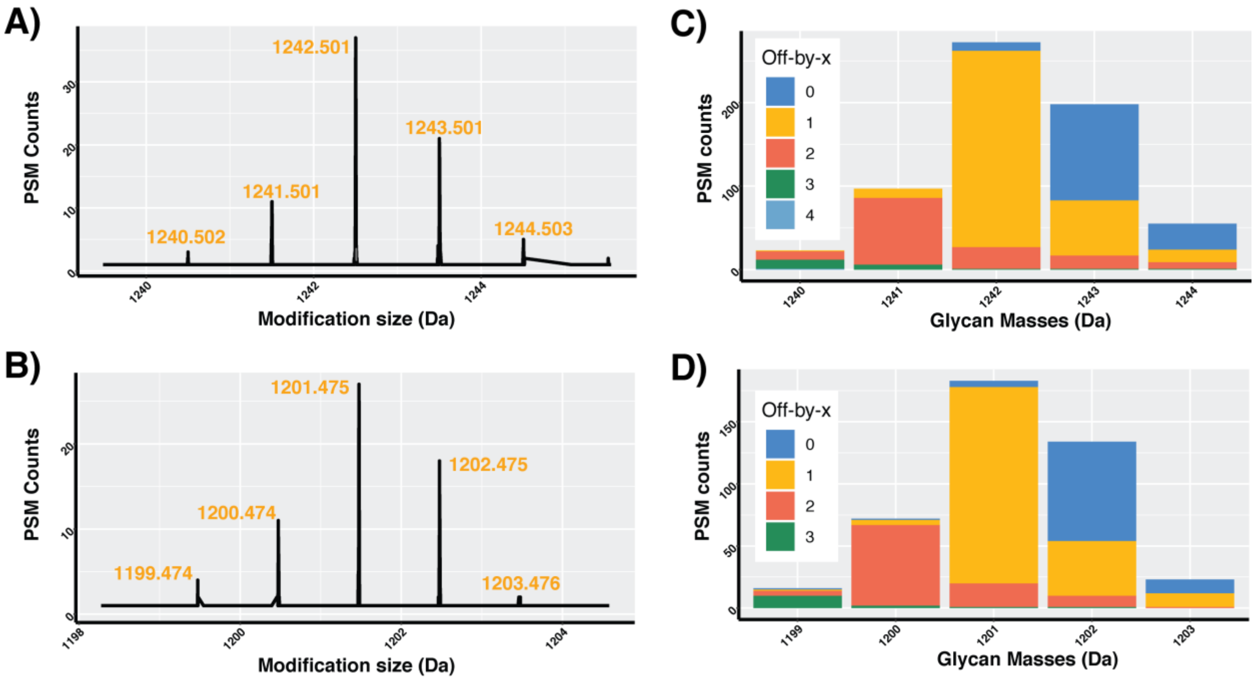


**Supplementary Figure 2: Detection of multiple delta masses associated with the expected *C. fetus fetus* glycans. A and B)** Observed delta masses binned in 0.001Da increments demonstrated clusters of PSMs separated by 1 Da characteristic of mis-assignment of the mono-isotopic mass. **C and D)** Examination of the "Off-by-x" parameter within Byonic demonstrates the majority of PSMs assigned as 1242 or 1201 have been corrected by one Dalton while the majority of PSMs assigned as 1243 and 1202 have had no corrections applied.

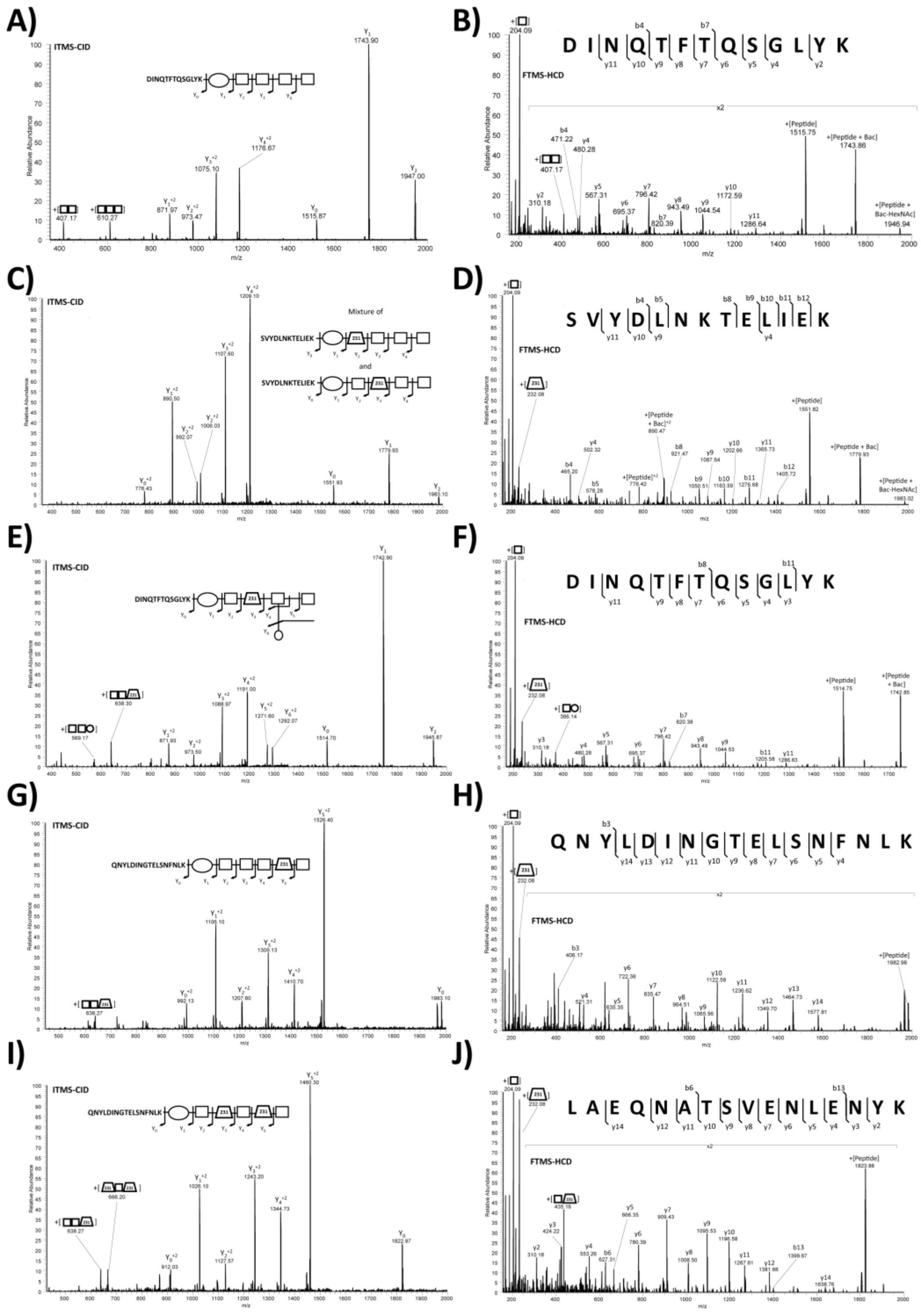


**Supplementary Figure 3: MS/MS analysis of novel *C. fetus fetus* glycoforms.** For each novel glycan the ITMS-CID and FTMS-HCD scans of a single precursor are provided confirming the presence of altered glycan structures. These annotations correspond to; **A and B)** HexNAc-HexNAc3-diNAcBac **C and D)** formylated HexNAc-HexNAc3-diNAcBac; **E and F)** formylated HexNAc-[Hex]-HexNAc3-diNAcBac; **G and H)** formylated HexNAc-[HexNAc]-HexNAc3-diNAcBac**; I and J)** double formylated HexNAc-[HexNAc]-HexNAc3-diNAcBac.

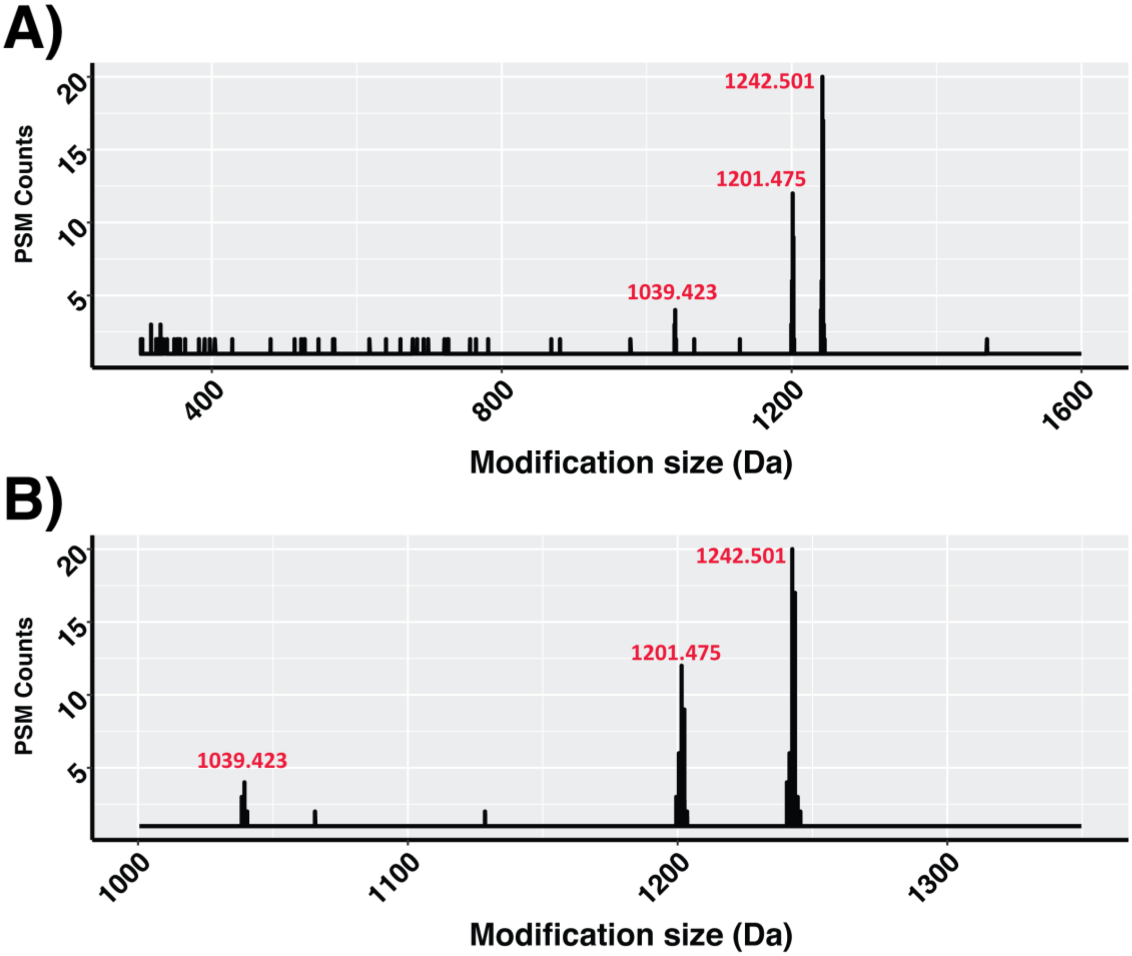


**Supplementary Figure 4: Absence of formylation on the glycans of unenriched *C. fetus fetus* glycopeptides.** Open searching of whole cell digests of *C. fetus fetus* demonstrates no formylation is observable prior to enrichment consistent with formylation being introduced during ZIC-HILIC enrichment.

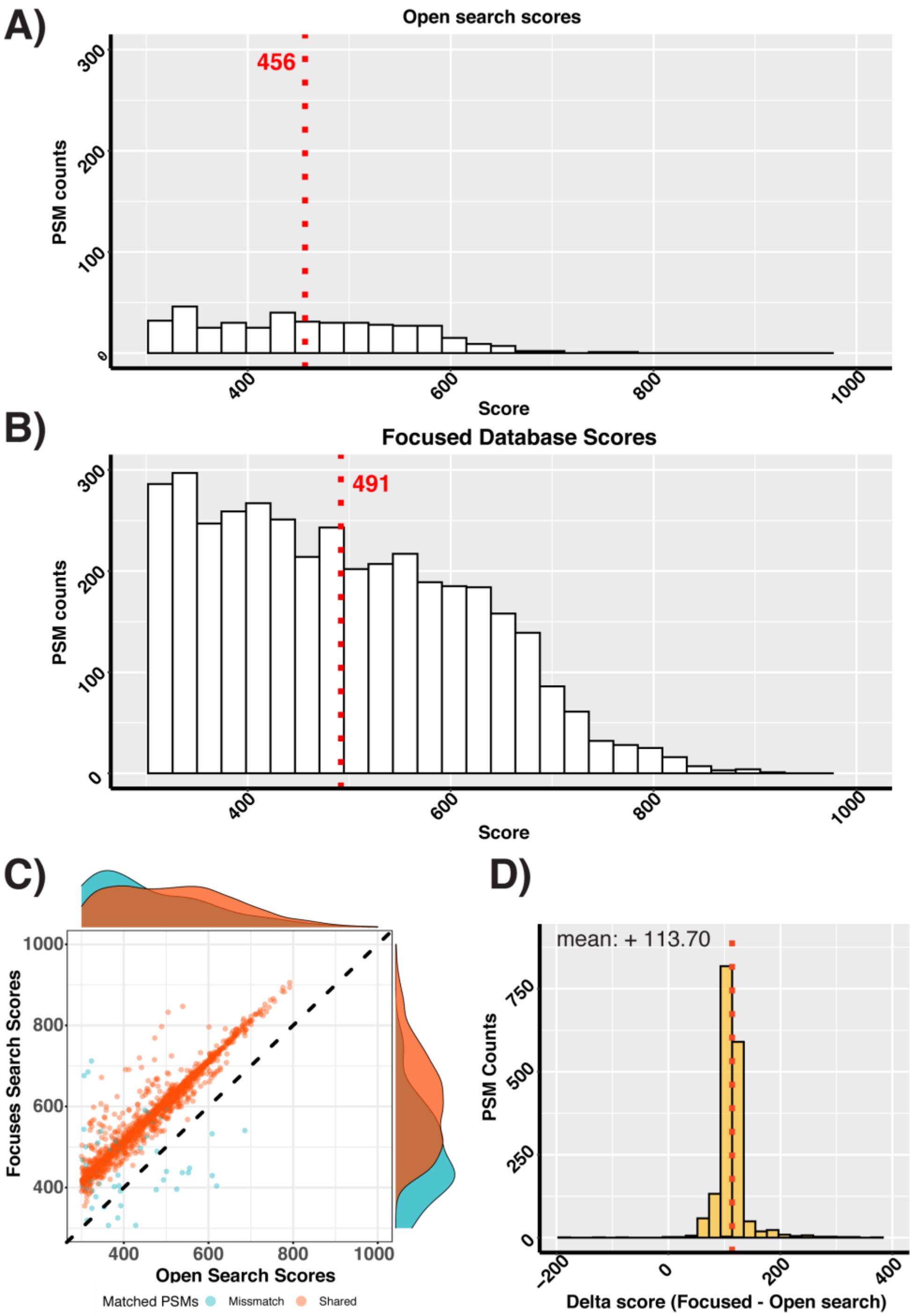


**Supplementary Figure: 5 Glycopeptide scores distributions observed using open and focused searches at the dataset and MS/MS scan levels.** For the seven glycans identified within *C. fetus fetus* (1039.423, 1067.419, 1201.475, 1229.469, 1242.501, 1270.497 and 1298.492) all glycopeptide PSMs are plotted for both Open (**A**) and focused searches (**B**). Comparison of glycopeptides scores for MS/MS scans identified within focused and open searches reveal the same spectra are typically assigned to the same peptide sequence but have a higher score in focused searches with a mean increase of ~113 (**C and D**)

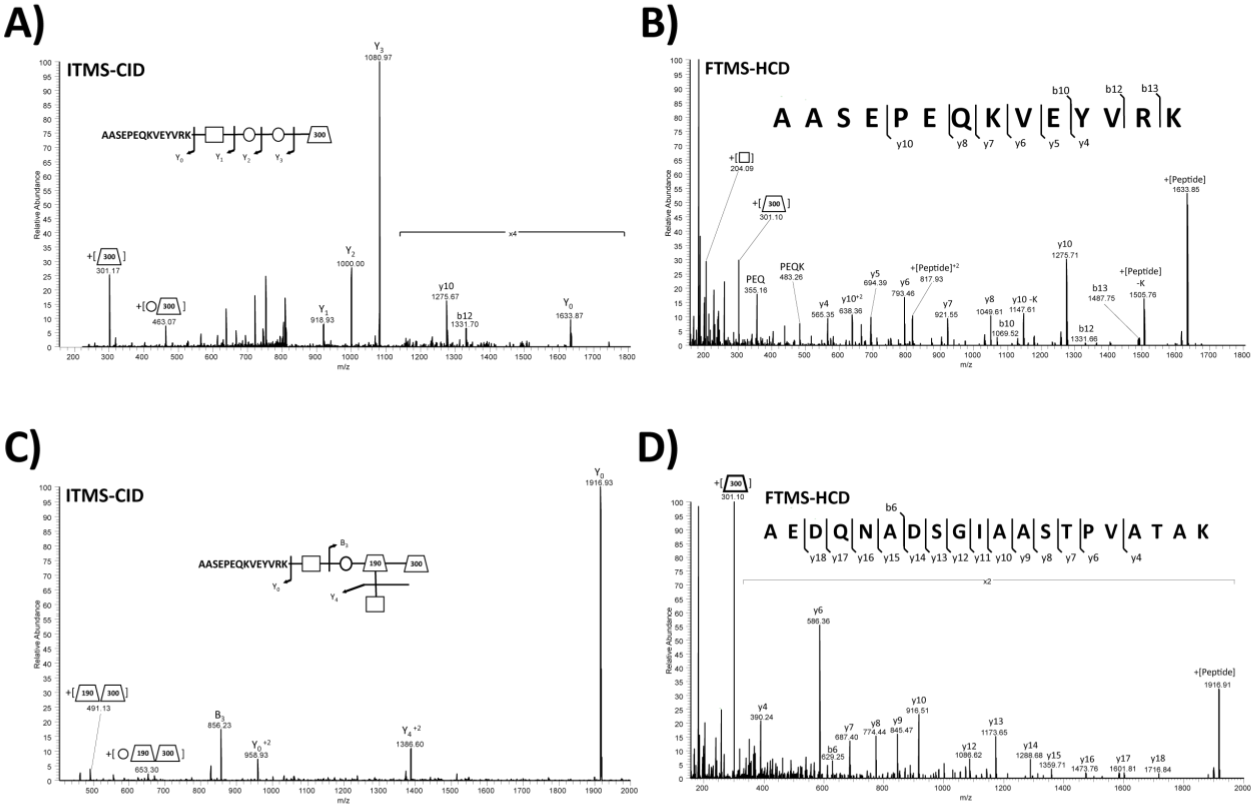


**Supplementary Figure 6: MS/MS analysis of novel *A. baumannii glycoforms.*** For each novel glycan the ITMS-CID and FTMS-HCD scans of a single precursor are provided confirming the presence of altered glycan structures. These annotations correspond to; **A and B)** HexNAc3NAcA4OAc-Hex2-HexNAc and **C and D)** formylated HexNAc3NAcA4OAc-[HexNAc]-Hex2-HexNAc.

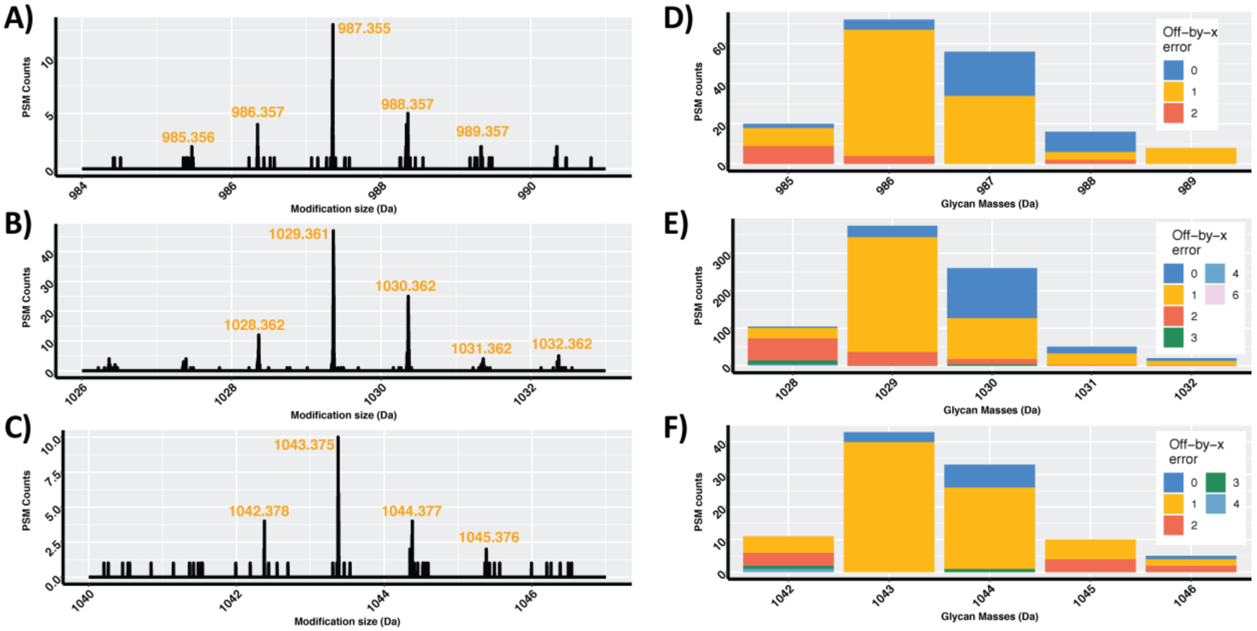


**Supplementary Figure 7: Detection of multiple delta masses associated with the expected *A. baumannii* glycans. A, B and C)** Delta masses binned in 0.001Da increments demonstrate clusters of PSM separated by 1 Da characteristic of mis-assignment of the mono-isotopic mass. **D, E and F)** Examination of the "Off-by-x" parameter within Byonic demonstrates the majority of PSMs assigned as 987, 1029 or 1043 have been corrected by one Dalton while a large proportion of PSMs assigned as 988, 1030 or 1044 have had no corrections applied.

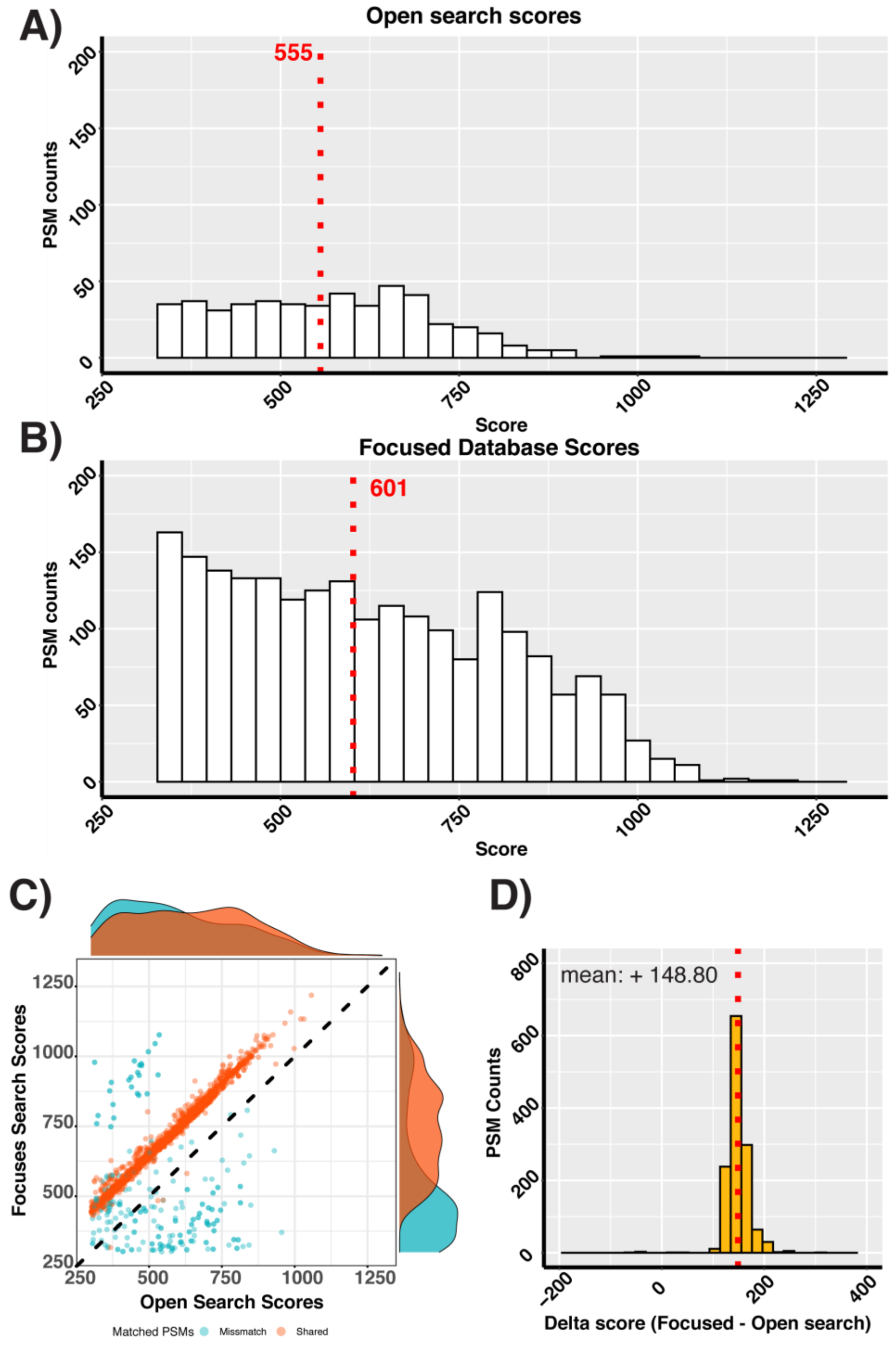


**Supplementary Figure 8 Glycopeptide scores distributions observed using open and focused searches at the dataset and MS/MS levels.** For the five glycans identified within *A. baumannii* (1030.368 Da, 988.357 Da, 1044.383 Da, 827.281 Da and 1058.358 Da) all glycopeptide PSMs are plotted for Open (**A**) and focused searches (**B**). Comparison of glycopeptides scores for MS/MS scans identified within focused and open searches reveal the same spectra are typically assigned to the same peptide sequence but have a higher score in focused searches with a mean increase of ~149 (**C and D**)

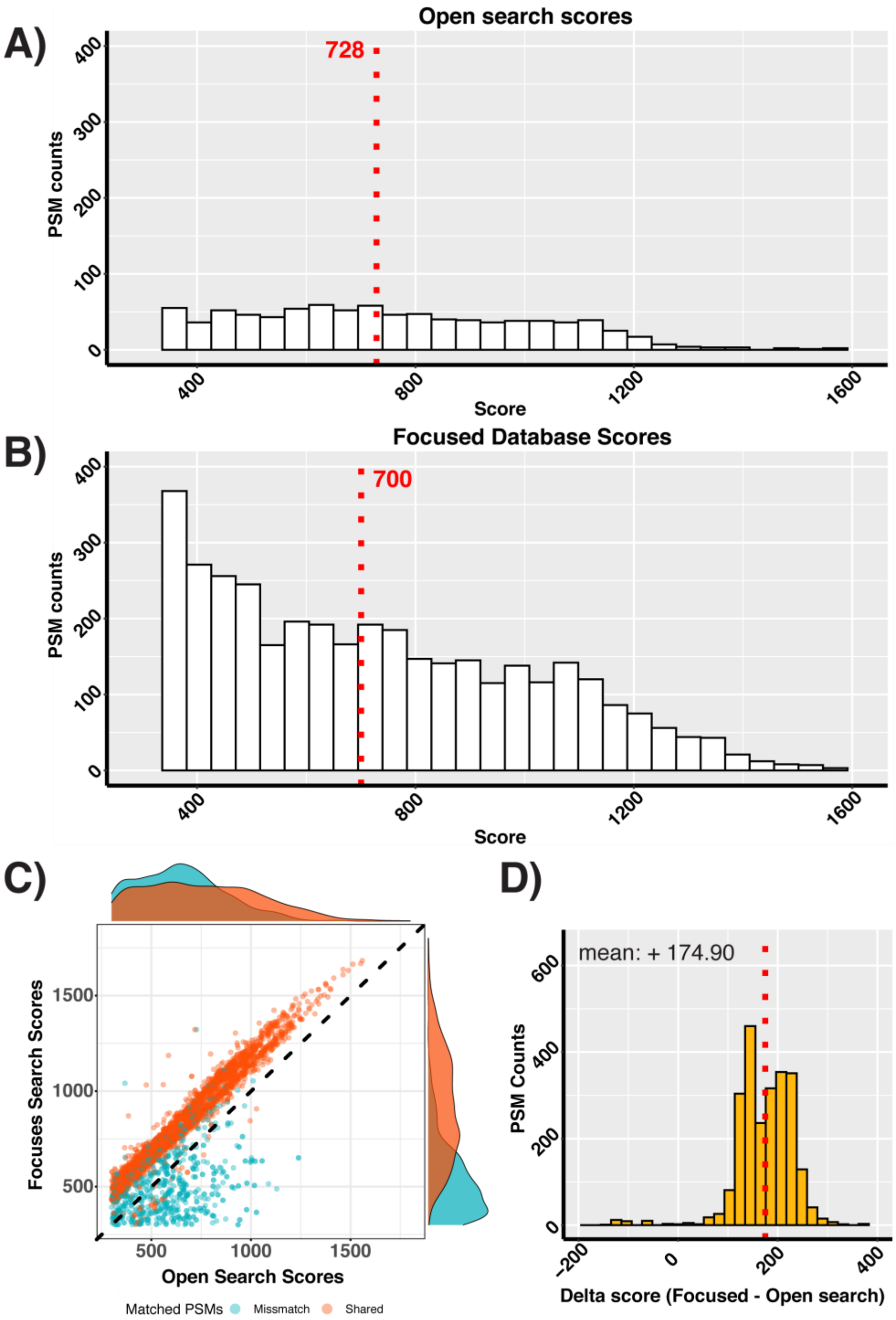


**Supplementary Figure 9 Glycopeptide scores distributions observed using open and focused searches at the dataset and MS/MS levels.** For the five glycans identified within *B. cenocepacia* (568.207Da, 696.202Da, 624.197, 668.223 and 696.218Da) all glycopeptide PSMs plotted for Open (**A**) and focused searches (**B**). Comparison of glycopeptides scores for MS/MS scans identified within focused and open searches reveal the same spectra are typically assigned to the same peptide sequence but have a higher score in focused searches with a mean increase of ~175 (**C and D**)

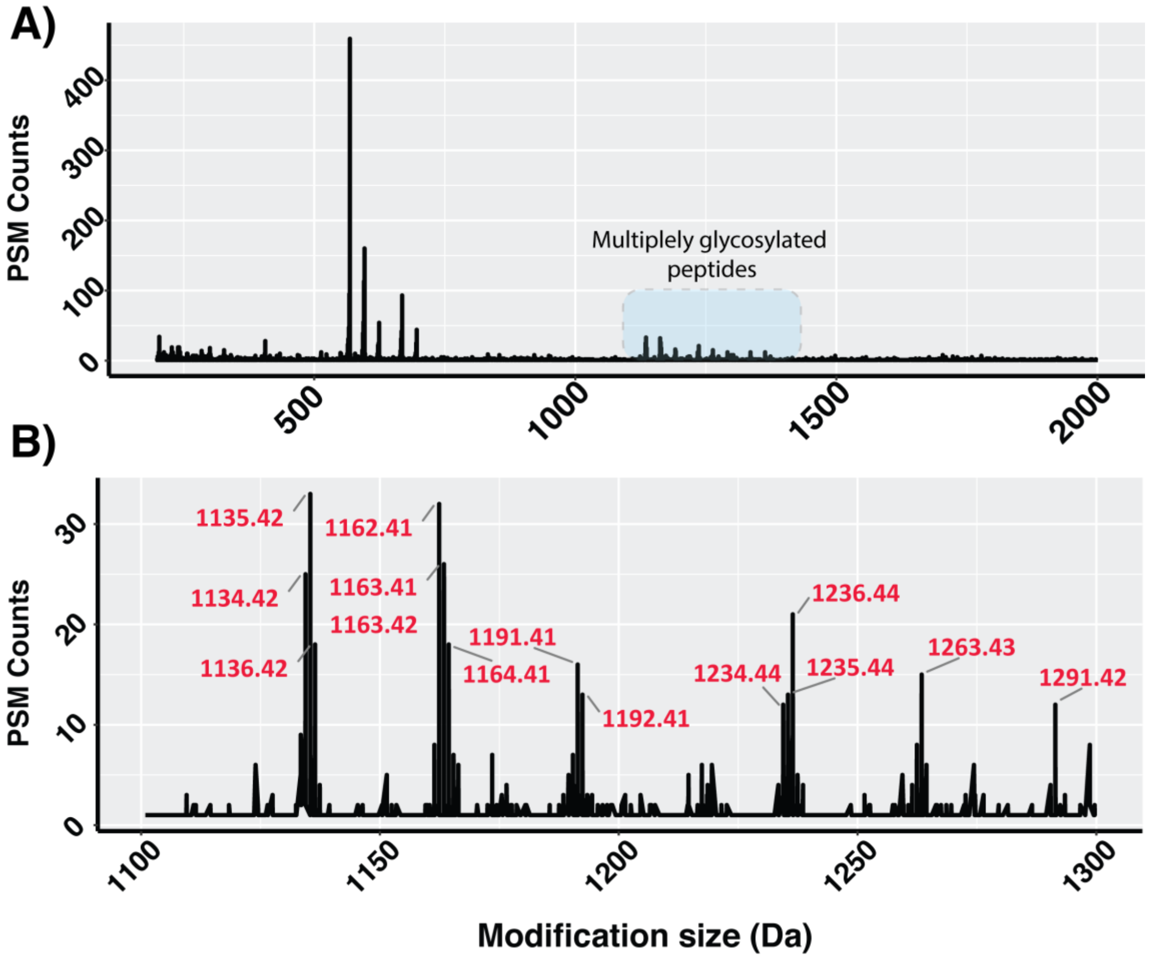


**Supplementary Figure 10: Open searches of *B. Cenocepacia* reveals masses consistent with multiply glycosylated peptides.** Examination of delta mass plots using 0.01Da mass bins demonstrates masses consistent with multiple glycans on assigned peptides. Note: 0.01Da mass bins have been used instead of 0.001 mass bins used in all other figures to more clearly highlight these delta masses.

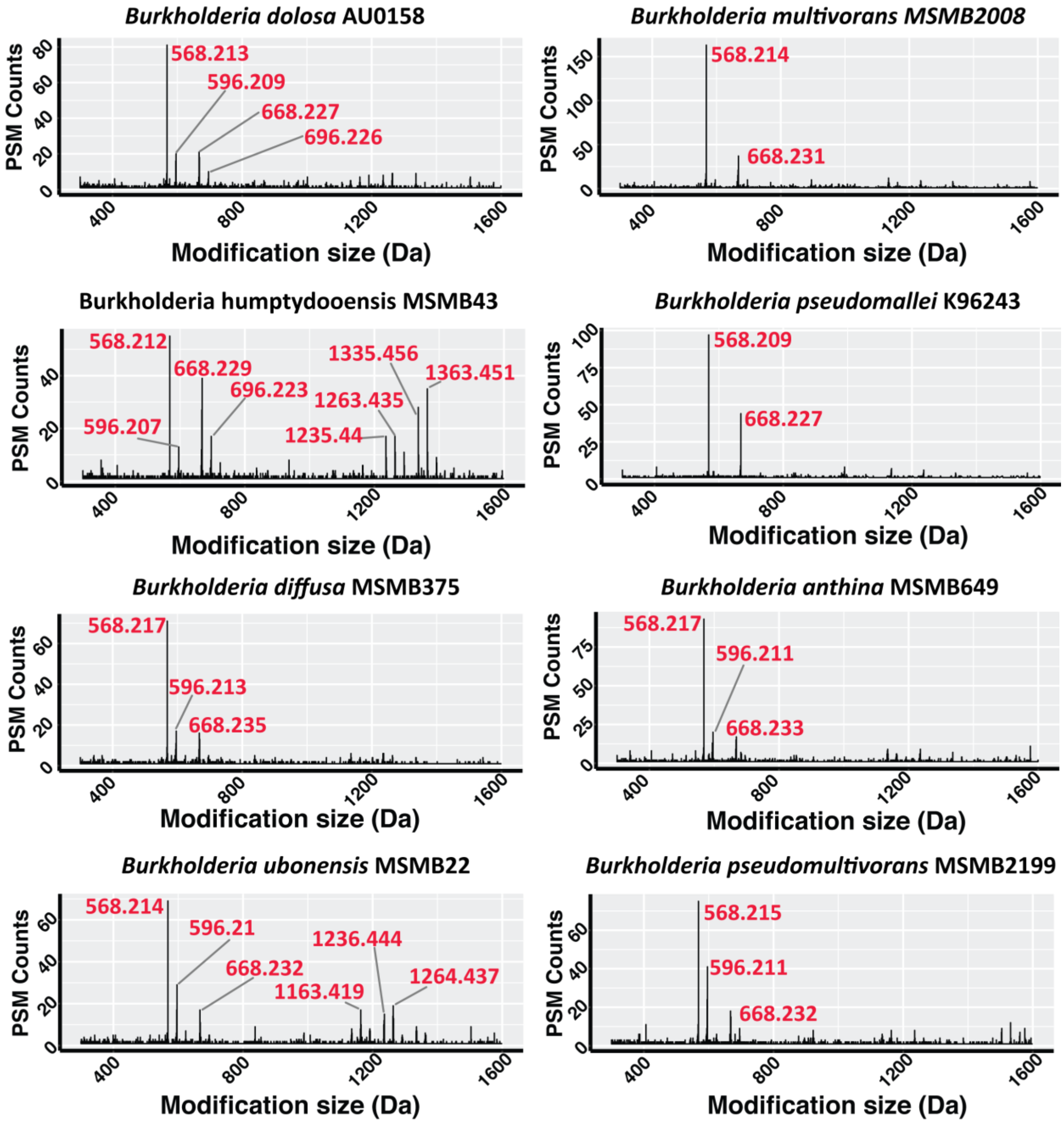


**Supplementary Figure 11: Delta mass profiles of Burkholderia species.** Examination of delta mass plots using 0.001Da mass bins demonstrates the presence of similar modifications across Burkholderia species.

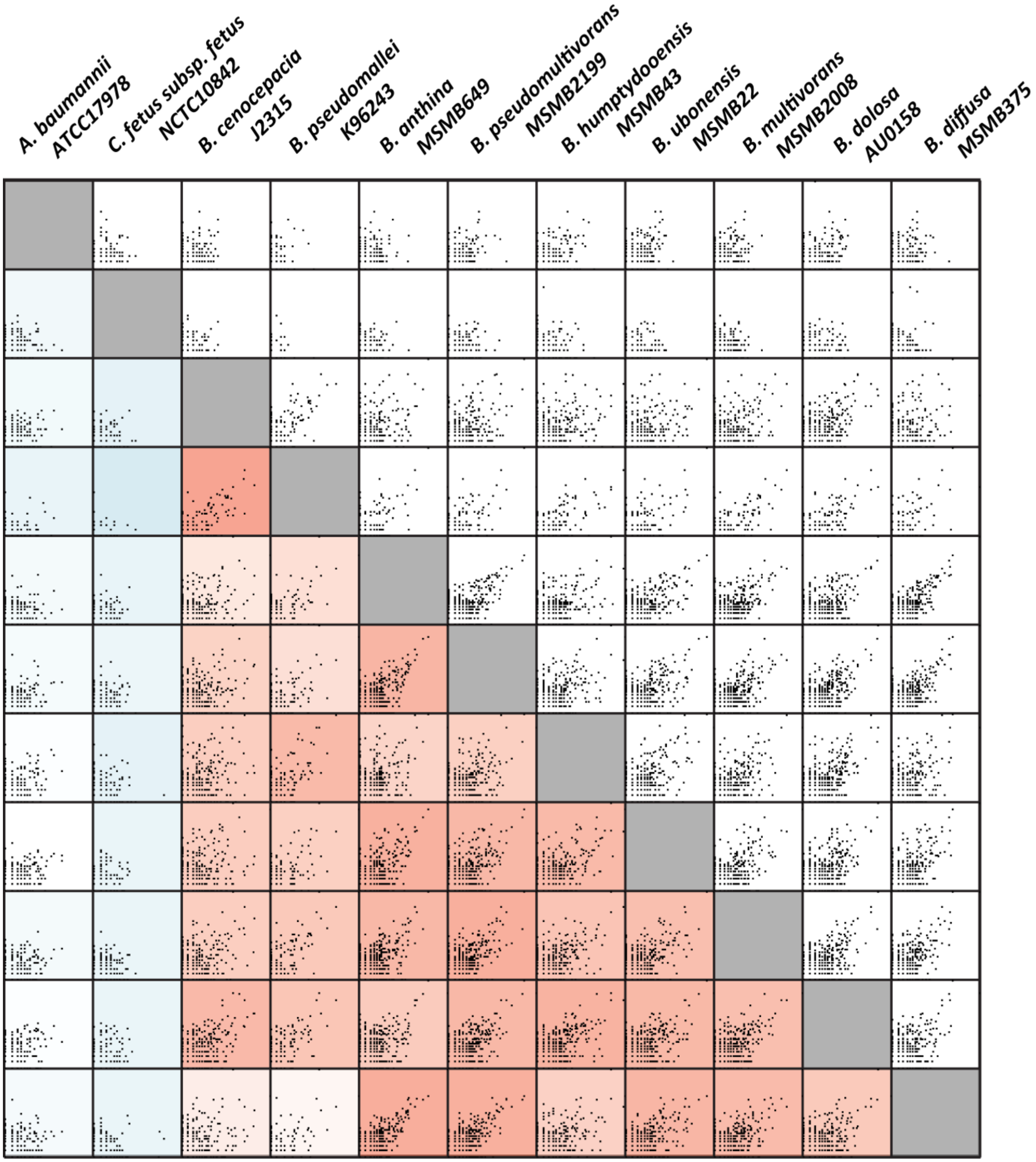


**Supplementary Figure 12: Pearson correlation analysis of delta mass profiles.** The Log2(PSM delta mass counts) of all delta mass profiles have been correlated using Pearson correlation and plotted.

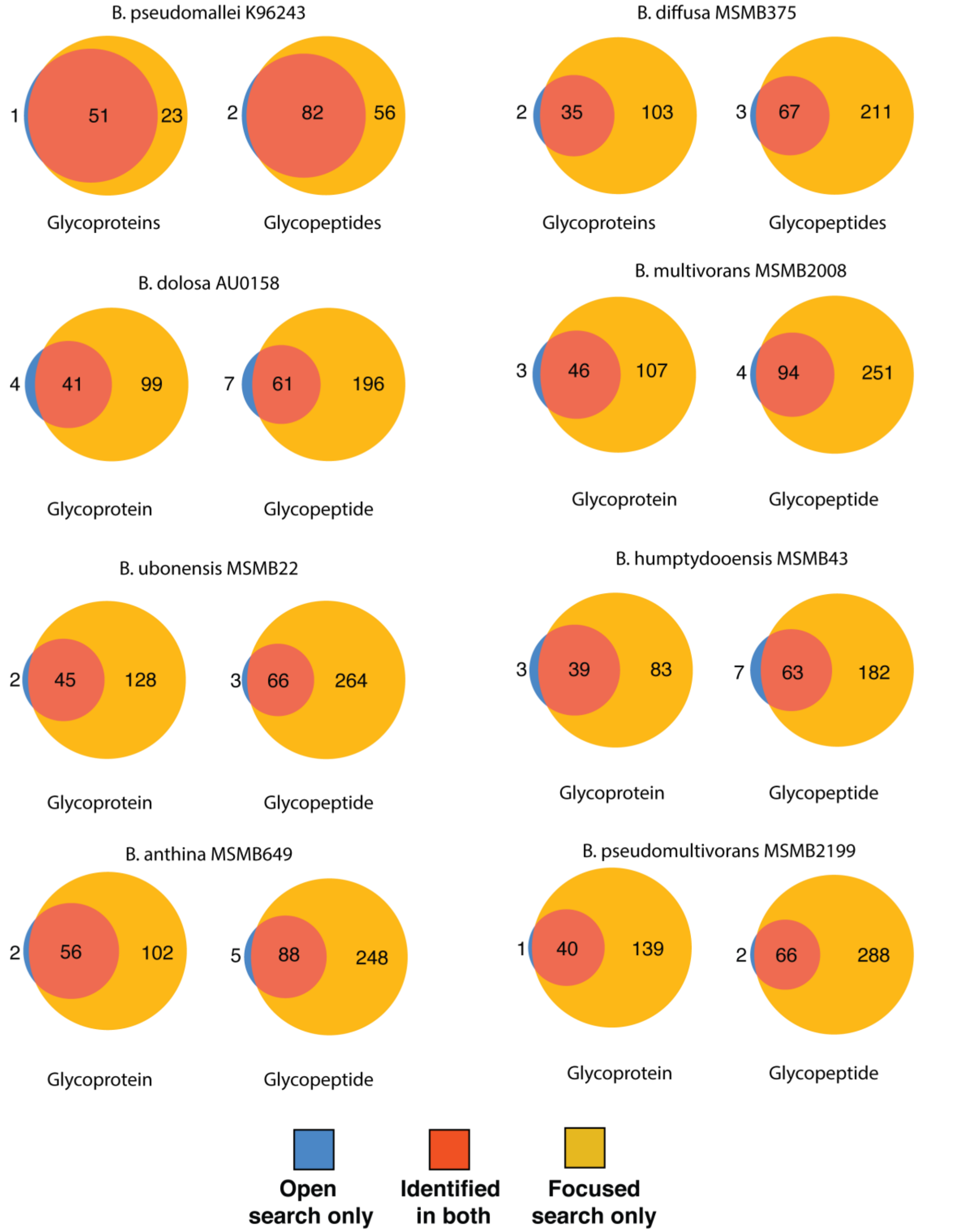


**Supplementary Figure 13: Comparison of the glycoproteome coverage between open and focused searches across Burkholderia species.** At the protein and peptide levels focused searches lead to a marked improvement in the observable glycoproteome.

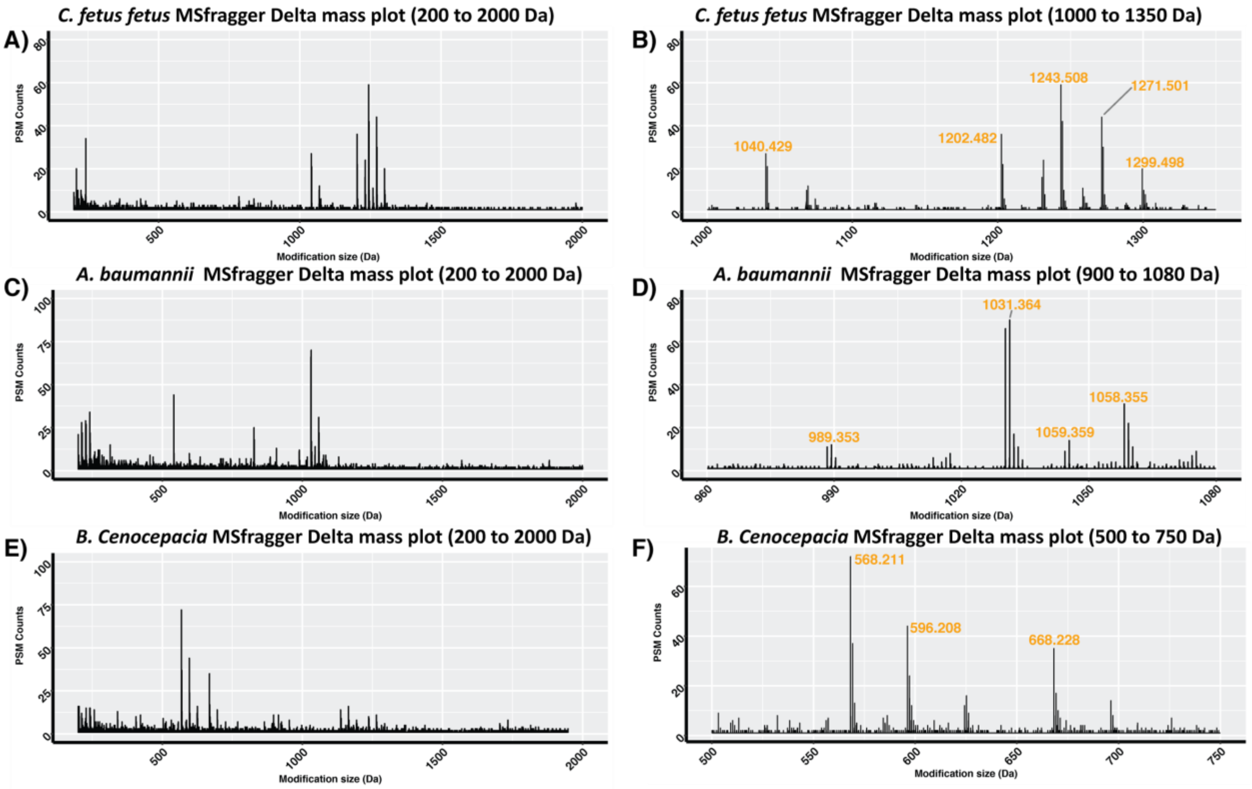


**Supplementary Figure 14: MSfragger based open searching of model strains.** Delta mass plots of the modification masses identified using MSfragger. Within glycopeptide enriched samples glycans of bacterial systems can be readily identified to a comparable level as Byonic open searches. **A and B)** *C. fetus fetus*; **C and D)** *A. baumannii* and **E and F)** *B. cenocepacia*.

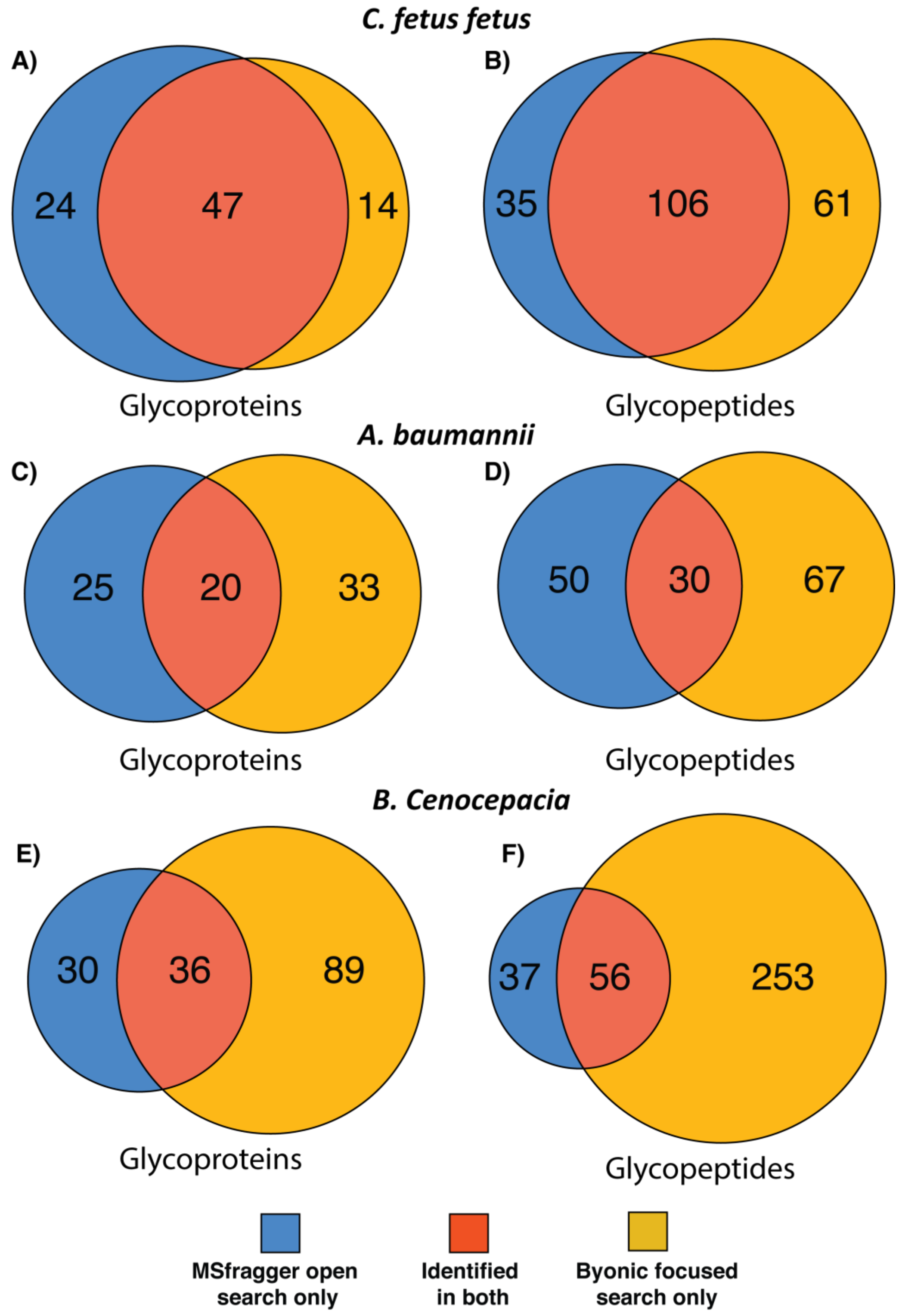


**Supplementary Figure 15: Comparison of glycopeptide and proteins identified between MSfragger searches and focused Byonic searches of model strains.** Venn diagrams showing the overlap between MSFragger results and the focused database searches in Byonic. **A and B)** *C. fetus fetus*; **C and D)** *A. baumannii* and **E and F)** *B. cenocepacia*.

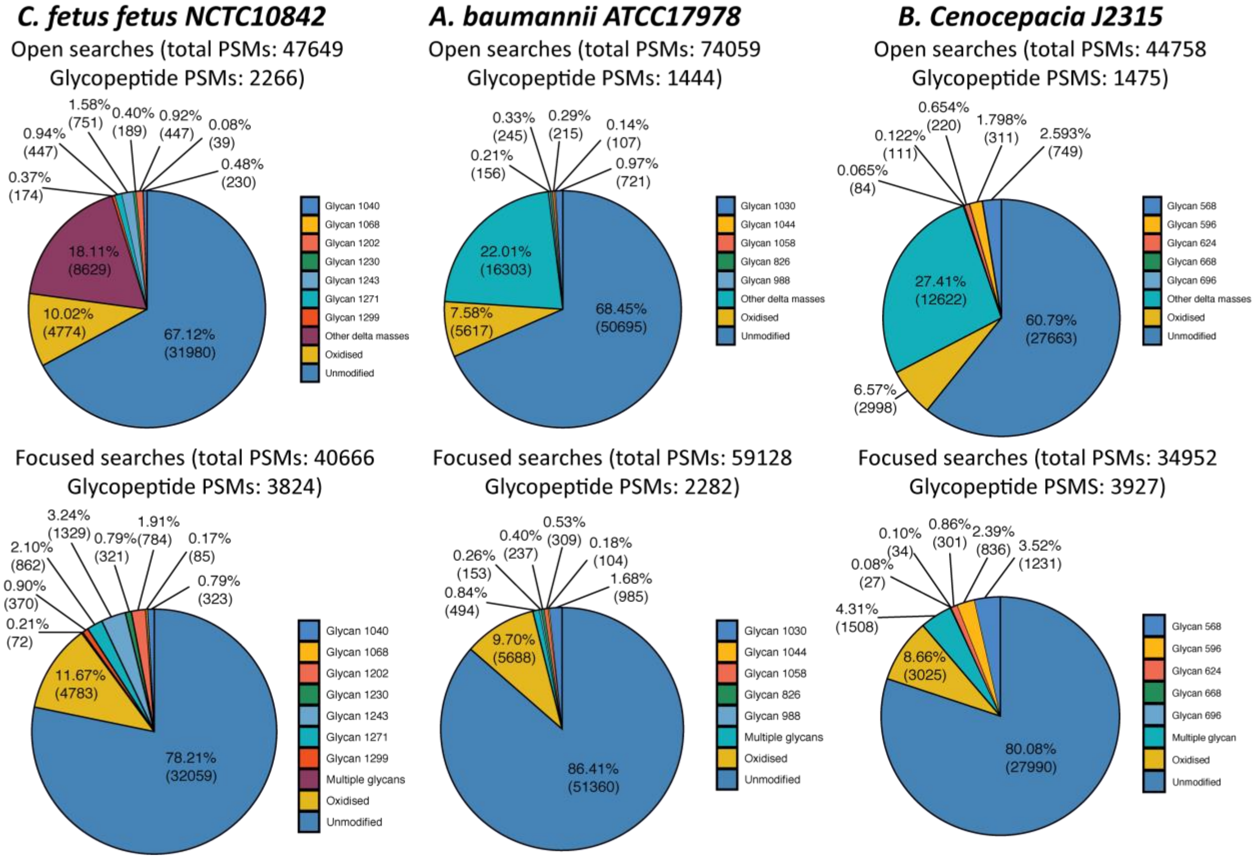


**Supplementary Figure 16: Comparison of glycopeptide PSMs identified using open searching or focused searches.** Glycopeptide PSMs account for <10% of all identified PSMs within ZIC-HILIC enriched samples.
